# Targeting NOTCH2-JAG1 juxtacrine signaling reverses macrophage-mediated tumor resistance to taxol

**DOI:** 10.1101/2024.07.08.602467

**Authors:** Fazhi Yu, Qin Zhou, Tong Zhou, Yijia Xie, Peng Zhang, Wei He, Weiqiang Yu, Aoxing Cheng, Hanyuan Liu, Qingfa Wu, Xiaopeng Ma, Jing Guo, Ying Zhou, Jue Shi, Zhenye Yang

## Abstract

Taxanes are widely used in chemotherapy, but primary and acquired resistance limit the clinical efficacy. Studies have shown tumor interaction with macrophages in the tumor microenvironment (TME) plays a significant role in taxane resistance, yet therapeutic strategies that directly deplete or repolarize macrophages are challenging and with considerable risk of side effects. Here we uncovered that tumor-macrophage interaction can be selectively targeted by inhibiting post-mitotic NOTCH2-JAG1 juxtacrine signaling in the TME, which strongly sensitizes paclitaxel response. Using translatome profiling, we found significant NOTCH2 upregulation during paclitaxel-induced prolonged mitosis. NOTCH2 was subsequently activated in the post-mitotic G1 phase by JAG1 expressed on the neighboring macrophages and tumor cells, which promoted tumor cell survival and upregulated cytokines that recruited JAG1-expressing macrophages, thus generating a positive feedback loop that further enhanced the pro-tumor NOTCH2 activity. By targeting this NOTCH2-JAG1 axis using NOTCH2 shRNA or a pan-NOTCH inhibitor, macrophage recruitment and paclitaxel resistance were significantly attenuated in multiple mouse tumor models of ovarian cancer. Clinical samples from paired primary and recurrent ovarian cancer patients also showed significant correlation of higher NOTCH2 expression with worse prognosis. Our results thus point to combining NOTCH2 inhibitor with taxane as an effective therapeutic strategy to selectively disrupt tumor-macrophage interaction in the TME and overcome macrophage-mediated taxane resistance in NOTCH2-positive tumors.

## Introduction

The anti-mitotic chemotherapeutics, taxanes, are employed as a major treatment modality for cancer patients, in particular those with solid tumors. However, primary and/or acquired resistances are common [1]. Much effort has been devoted to developing combinatorial strategies that can sensitize tumor response to taxanes for the resistant tumor types. A key bottleneck of developing such combinatorial strategy is in identifying tumor-selective targets. Many common tumor intrinsic targets, such as kinases and immuno-oncology targets (e.g., immune checkpoints and cytokines), have essential functions in normal physiology so drugging them inevitably increases the risk of toxicity and systemic side effects in essential organs [2]. One possible avenue to develop more tumor-selective combinatorial strategies to sensitize taxane response is to explore and target chemoresistance-associated signaling that are highly activated only in the tumor microenvironment (TME), e.g., via cell-cell interaction in the TME.

Taxanes, i.e., paclitaxel and its derivatives, bind to the β tubulin and block microtubule dynamics [3, 4]. This results in aberrant spindles and erroneous microtubule-kinetochore attachments during mitosis, leading to activation of the spindle assembly checkpoint (SAC) and prolonged mitotic arrest [5]. Taxane-treated cells may die in the mitotic arrest, undergo multipolar division, or slip out of mitosis without division to an abnormal G1 state, from which the cells may die, arrest or progress through cell cycle [6–12]. Taxane resistance has mostly been characterized for tumor intrinsic mechanisms, such as upregulated drug efflux, activation of pro-survival pathways and/or inactivation of apoptotic pathways, altered tubulin expression or dynamics, and metabolic alterations [13–16]. Recently the involvement of TME in taxane resistance has been reported, e.g., via cytokines secreted by cancer-associated fiboblasts and adipocytes [17, 18]. A number of *in vivo* studies also showed that immune cells residing in the TME contribute to taxane resistance [19–23]. Particularly, tumor-infiltrated macrophages were found to play a crucial role in conferring resistance to taxol-induced growth arrest and cell death, and systemic depletion or repolarization of macrophages can reverse the resistance [19, 20, 24–26]. Nonetheless, we still lack mechanistic understanding of the specific contact-dependent signals that link tumor cells with other cell types in the TME and subsequently promote chemoresistance.

To uncover novel cell-cell interactions in the TME that are specific to paclitaxel resistance and may be selectively targeted, in this study we combined translatome profiling of paclitaxel-induced mitotic cells as well as co-culture analysis of cancer cells and macrophages. We found NOTCH2 receptor in cancer cells was selectively upregulated during taxol-induced prolonged mitosis. Interaction of NOTCH2 with JAG1 mainly expressed on the neighboring macrophages activated juxtacrine signaling in the post-mitotic G1 state, which promoted resistance to taxol-induced cell death. Activation of NOTCH2 signaling via tumor-macrophage interaction also upregulated cytokines that stimulated the recruitment of JAG1-expressing, suppressive macrophages, further enhancing taxol resistance. Our results thus suggested NOTCH2 is a viable combinatorial target to selectively attenuate tumor-macrophage interaction in the TME and paclitaxel resistance. Indeed knocking down NOTCH2 by shRNA or using a pan-NOTCH inhibitor, RO4929097, in combination with paclitaxel, was able to significantly suppress tumor growth in multiple mouse tumor models of ovarian cancer that were resistant to paclitaxel treatment alone, illustrating that NOTCH2 inhibitor is an effective combinatorial drug candidate to target tumor-macrophage interaction specifically activated in the TME to reverse taxol resistance.

## Results

### Taxol induces translational upregulation of the transmembrane receptor NOTCH2

To identify cell surface receptor(s) that mediates cell-cell interaction in the TME and potentially contributes to taxol resistance, we first conducted translatome profiling of upregulated membrane proteins/receptors in taxol-resistant cancer cells during drug-induced prolonged mitotic arrest. We hypothesized that as prolonged mitosis is the primary phenotype of taxol, cellular alternations occurring during the mitotic arrest likely play crucial roles in engendering chemoresistance. And since transcription is largely silenced during mitosis, upregulation of pro-tumor molecular mediators probably occurs at the post-transcriptional level, such as translation. To perform the mitotic translatome analysis, mitotic cells were harvested by shake-off from untreated and taxol-treated HeLa cells and subjected for ribosome profiling (Figure 1A). More than nine thousands mRNAs were identified to be translated during normal mitosis, and 89% of these genes were also found in two previously published datasets, in which the translational landscape of mitosis was analyzed also by ribosome profiling [27, 28], validating that our data are well in line with results from previous studies (Figure S1A). From the genes identified to be upregulated during prolonged mitosis (Figure 1A), cell membrane receptors were picked out and compared with those identified from two other available translatome analysis of prolonged mitosis using ribosome profiling [27] and proteomics [29], respectively. The Venn diagram showed that NOTCH2 is the only upregulated membrane receptor identified by all three studies (Figure 1B).

**Figure 1.**
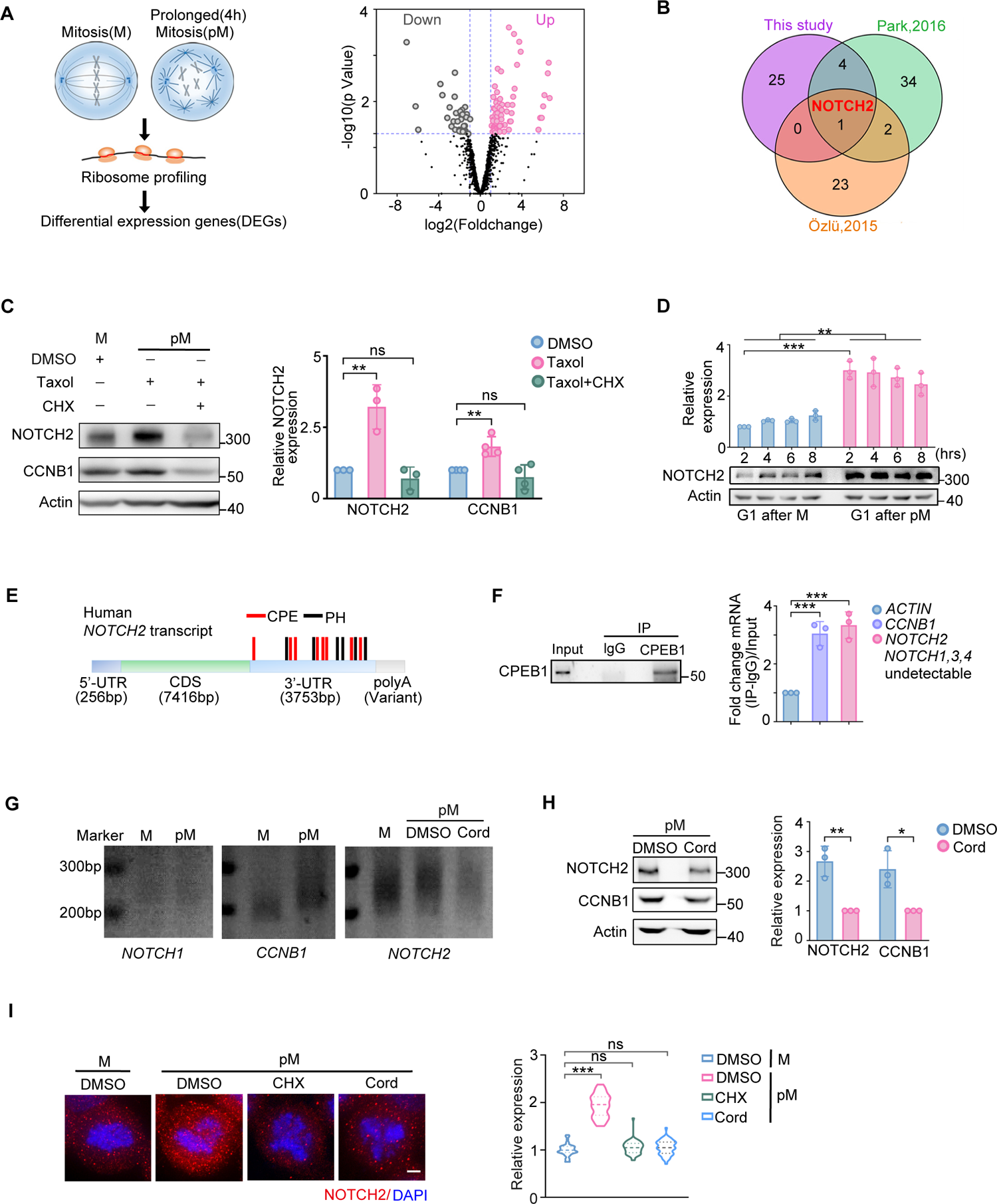
NOTCH2 translation is upregulated during taxol-induced prolonged mitosis. **(A)** The diagram showed the ribosome profiling process and the Volcano-plot of mRNA expressions of prolonged mitotic (pM) versus mitotic (M) cells. Significantly upregulated genes (log2 (Fold change)>1.8 and pValue<0.05) and downregulated genes (log2 (Fold change) <-1.8 and pValue<0.05) were presented. **(B)** Venn diagram showing the differentially expressed genes (DEGs) identified by ribosome profiling conducted in this study and Park, 2016 [27], and the genes that expressed higher on the membrane of mitotic cells than interphase cells from results by Özlü, 2015 [29]. **(C)** Western blot analysis of NOTCH2 and cyclin B1 (CCNB1) in HeLa cells in mitosis (M) and prolonged mitosis (pM) induced by taxol. Cells in prolonged mitosis were further treated with DMSO or 10 μM Cycloheximide (CHX) for 4 hours. Quantified results are shown on the right and error bars are SEM. **(D)** HeLa cells in mitosis (M) and prolonged mitosis (pM) induced by 0.5 μM Nocodazole were harvested by shake-off and re-plated in a culture dish with fresh medium for various hours before being collected. Western blot was conducted to analyze the protein levels of NOTCH2. **(E)** Schematic representation of the number and position of CPE elements, polyadenylation hexanucleotide (PH) and polyadenylation signals in the 3’-UTR of the Human NOTCH2 transcript. **(F)** RT–PCR detection of mRNA levels of the indicated genes after immunoprecipitation of CPEB1 in HeLa cells in prolonged mitosis. The CPEB1 IP efficiency was confirmed using western blot. **(G)** Poly(A) tail-length assay of NOTCH1, CCNB1 and NOTCH2 transcripts of HeLa cells in mitosis (M) and prolonged mitosis (pM). Cells in prolonged mitosis were treated with DMSO or 10 μM Cordycepin (Cord, CPE inhibitor) for 4 hrs before collection. **(H)** Western blot analysis of NOTCH2 and cyclin B1 in HeLa cells in prolonged mitosis (pM) (left). Cells in prolonged mitosis were treated with DMSO or 10 μM Cordycepin (Cord, CPE inhibitor) for 4 hrs before collection. Quantified results are shown in the right panel. (**I**) Immunofluorescence was used to detect the expression levels of NOTCH2 in HeLa cells in mitosis (M) and prolonged mitosis (pM) induced by taxol, followed by quantitative analysis. Data are shown as mean ± SD, and data of (C), (D), (F), (H) and (I) were averaged from at least three replicates of each sample and/or treatment condition. p values were determined by two-tailed student’s t test (ns, not significant; *p < 0.05, **p < 0.01, ***p < 0.001).

NOTCH2 is one of the four NOTCH signaling receptors that are frequently dis-regulated in cancers and involved in multiple oncogenic phenotypes, such as drug resistance via juxtacrine signaling at cell-cell contact [30–32]. We first validated NOTCH2 was upregulated at the protein level during mitotic arrest. Similar to cyclin B, which is known to be actively translated during prolonged mitosis [15], the level of NOTCH2 protein (Figure 1C), but not the mRNA (Figure S1B), significantly increased during taxol-induced prolonged mitosis. The increase of NOTCH2 protein was abolished by cycloheximide treatment (Figure 1C), indicating that NOTCH2 was upregulated at the protein synthesis level. While the protein level of NOTCH2 was already at a high level when cells entered the G1 phase (Figure 1D), the mRNA of NOTCH2 was not increased until six hours after mitotic exit (Figure S1C), supporting the translational upregulation of NOTCH2 during prolonged mitosis. We also confirmed the upregulation of NOTCH2 protein in two other cancer cell lines, the ovarian cancer line OVCAR8 and the lung cancer line A549 (Figure S1D). In addition, we found NOTCH2 upregulation can be induced by other anti-mitotic compounds, such as nocodazole, STLC (Eg5 inhibitor) and BI2536 (Plk1 inhibitor) (Figure S1E). Overall our results showed that NOTCH2 is actively translated and upregulated during taxol-induced prolonged mitosis.

### CPEB-mediated polyadenylation promotes translation of NOTCH2 during prolonged mitosis

Uniquely among the four NOTCH receptors, NOTCH2 is the only one upregulated during prolonged mitosis (Figure S1F), despite that both NOTCH2 and NOTCH3 are expressed in ovarian and lung cancers according to the TCGA datasets (Figure S1G). To explore the origin of the differential translation of distinctive NOTCH receptors in prolonged mitosis, we examined the characteristics of NOTCH mRNA in comparison with the mRNA of cyclin B, a highly translated protein in mitosis. Cyclin B mRNA has multiple cytoplasmic polyadenylation element (CPE) and polyadenylation hexanucleotide (PH) elements. It has been reported that CPEB (cytoplasmic polyadenylation element binding protein) and CPSF (cleavage and polyadenylation specificity factor) promote the polyadenylation and translation of mRNA that contain sequences of CPE and PH [33]. Similar to cyclin B, there are eight CPE and five PH elements within the 3’ UTR of NOTCH2 mRNA (Figure 1E and Figure S1H). In contrast, few CPE or PH sequences were found in the mRNA of the other three NOTCH receptors (Figure S1H). CPEB RIP assay demonstrated that similar to cyclin B, whose mRNA is known to bind CPEB, mRNA of NOTCH2 also bound to CPEB1 in the mitotic cells (Figure 1F), suggesting possible polyadenylation of NOTCH2 mRNA during prolonged mitosis. We then analyzed the status of the polyA tail within the mRNA by PCR. Similar to cyclin B, NOTCH2 mRNA showed higher polyadenylation during prolonged mitosis, whereas NOTCH1 mRNA did not showed polyadenylation (Figure 1G). In addition, CPEB inhibition with cordycepin, an inhibitor of CPEB-dependent polyadenylation, significantly attenuated NOTCH2 upregulation (Figure 1H, I), illustrating that CPEB-directed polyadenylation promoted the synthesis of NOTCH2 during prolonged mitosis.

### NOTCH2 activation confers taxol resistance

NOTCH and the downstream signaling cascades are activated by NOTCH binding to its cognate ligands expressed on the neighboring cells. Well-known NOTCH ligands include JAG1, JAG2, DLL1, DLL3 and DLL4 [31]. Based on analysis of the TCGA database of these ligands, we found JAG1 exhibited high expression in lung and ovarian cancer at both the mRNA (Figure 2A) and protein level (Figure S2A). To investigate the involvement of NOTCH2 activation in taxol response, we first used recombinant JAG1 that was pre-coated on coverslip to activate NOTCH2 in A549 cells, which indeed resulted in nuclear accumulation of NOTCH2 and HES1 (a known NOTCH2 target) (Figure 2B). Further live cell imaging analysis showed that pre-coated FC-rJAG1, but not FC, significantly reduced cell death in the post-mitotic G1 phase under taxol treatment, and this pro-survival phenotype was abolished in NOTCH2 knockout cells (Figure 2C, Figure S2B). Moreover, overexpression of the active intracellular domain of NOTCH2 (NICD2) downregulated the active caspase-3 level in taxol-treated cells (Figure 2D) and promoted tumor growth in mice treated with traxol (Figure 2E). Overall both the *in vitro* and *in vivo* results suggested that upregulated NOTCH2 signaling via interacting with JAG1 promotes tumor resistance to paclitaxel treatment.

**Figure 2.**
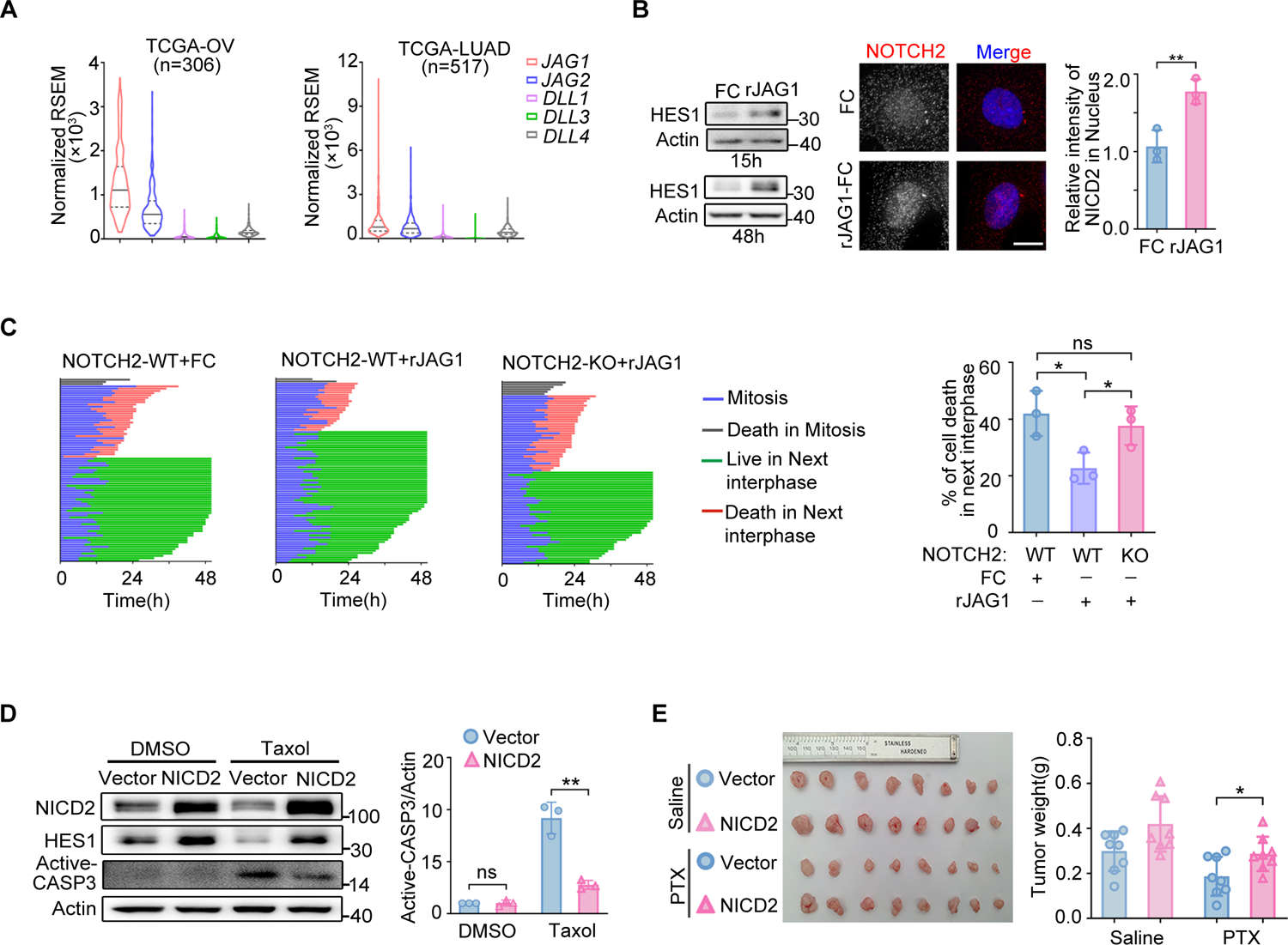
Activation of NOTCH2 signaling confers resistance to taxol-induced cell death. **(A)** Analysis of mRNA levels of NOTCH ligands in ovarian and lung cancer using TCGA data. **(B)** Western blot analysis of NOTCH signaling activity by HES1 expression in A549 cells plated on Fc or JAG-1 Fc for 15 hrs or 48 hrs. Immunofluorescence detection of NOTCH2 receptor in A549 cells. A549 cells were plated on Fc or JAG-1 Fc for 15 hrs, and NOTCH2 was analyzed using immunofluorescence. Scale bar, 10 μm. Quantification of NICD2 fluorescence intensity in cell nucleus is shown using relative fluorescence units. **(C)** Cell fate profiles of A549 cells with NOTCH2 KO in comparison with WT upon interacting with Fc or JAG-1 Fc. A549 cells were treated with Fc or JAG-1 Fc for 15 hrs before taxol treatment and then imaged by time-lapse microscopy under taxol for 72 hrs. **(D)** Western blot analysis of NICD2, HES1, active Caspase-3 and loading control in A549 cells with overexpression of the activated intracellular domain of NOTCH2 (NICD2). **(E)** Representative dissected tumors and tumor weight of xenograft mice with control A549 cells or A549 cells over-expressing the activated intracellular domain of NOTCH2 (NICD2). Data are shown as mean ± SD, and data of (B) and (C) were obtained from at least three experimental replicates. p values were determined by two-tailed Student’s t test (ns, not significant; *p < 0.05, **p < 0.01).

### JAG1 expressed on the neighboring macrophages is primarily responsible for activating NOTCH2 and promotes taxol resistance

By performing immunostaining (Figure 3A, B) and immunochemistry (Figure S3A) of ovarian tumor section and ascites from patients (Figure 3B), we confirmed the expression of JAG1 in ovarian tumor cells, and also found JAG1 was even more highly expressed in CD68^+^ macrophages in the TME (Figure 3A, B). Further flow cytometry analysis confirmed that while JAG1 was expressed in human ovarian cancer cell lines (Figure S3B), it was expressed at a significantly higher level in macrophages from the ascites of human ovarian tumors treated with taxol (Figure 3C). These data suggested JAG1 expressed on the neighboring macrophages is potentially the major activator of *in vivo* NOTCH2 signaling in the TME that underlies taxol resistance.

**Figure 3.**
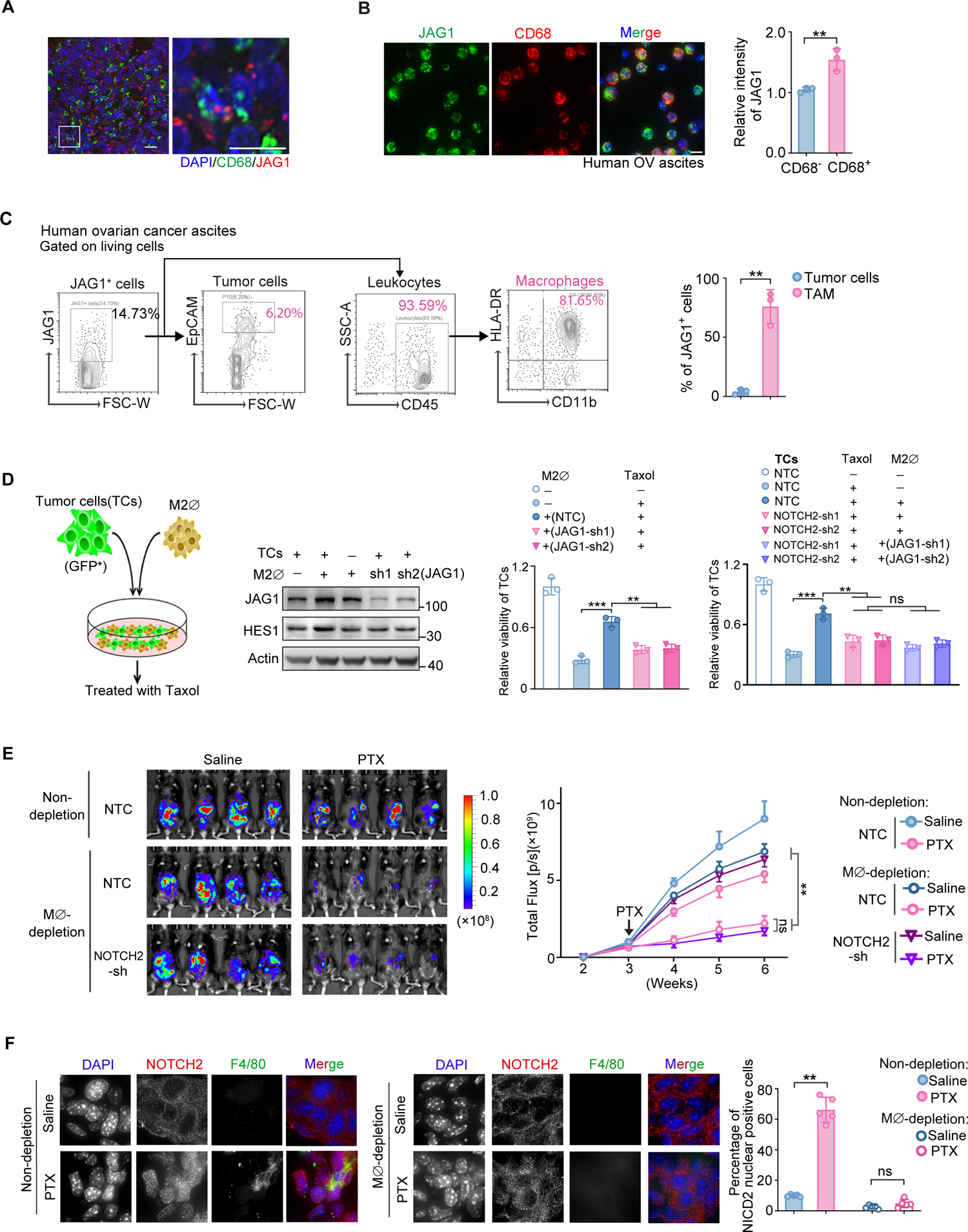
JAG1 expressed on the macrophages is primarily responsible for activating NOTCH2 signaling and promotes tumor resistance to taxol. **(A)** Immunofluorescence of DNA (blue), JAG1 (red) and macrophages marker (CD68, green) in tumor section from recurrent ovarian cancer patients. Framed image was enlarged on the right. **(B)** Immunofluorescence of JAG1 expressed on macrophages in the ascites of ovarian cancer patients. Representative images and quantification of relative intensity of JAG1 are shown. Three ascites were analyzed for each group. Scale bars, 10 μm. **(C)** FACS analysis of the ratios of tumor cells and macrophages among the JAG1^+^ cells from the ovarian cancer patient ascites. **(D)** Co-culture assay of OVCAR8 cells with THP-1-derived M2-macrophage (M2ϕ). The diagram shows the experimental set-up (left panel). Western blot analysis of JAG1, HES1 and the loading control for OVCAR8 cells in co-culture with THP-1-derived M2-macrophage (M2ϕ) (middle left panel). THP-1 cells were transduced with two different shRNA clones of JAG1 or vector control, then induced to be M2 macrophage M2 (M2ϕ) and co-cultured with OVCAR8-GFP cells in the presence of DMSO or paclitaxel (middle right panel). The OVCAR8-GFP cells were transduced with two different shRNA clones of NOTCH2 or vector control before co-cultured with THP-1-derived M2-macrophage cells (right panel). Quantification of the relative viability of GFP^+^ cells is shown. **(E)** Representative images of mice following the depletion of macrophages using liposome-clodronate that carried ID8-luciferase cells (left panel). The corresponding tumor growth curves are displayed in the right panel. **(F)** Immunofluorescence of NICD2-positive cells in tumor tissues from the ID8 mouse model and the quantified results. Data are shown as mean ±SD. Data of (B), (C) and (D) were obtained from three experimental replicates. p values were determined by two-tailed Student’s t test (**p < 0.01, ***p < 0.001).

To confirm and explore the role of JAG1-expressing macrophages in the observed taxol resistance, we used both *in vitro* co-culture models of OVCAR8 cells with M2-type macrophages derived from THP-1 (a human leukemic monocyte cell line, Figure 3D) as well as *in vivo* mouse tumor models with macrophage depletion (Figure 3E, F). We found that THP-1-derived M2 macrophages had a higher JAG1 expression than the cancer cells (Figure 3D). While the presence of M2 macrophages significantly promoted the survival of OVCAR8 cancer cells, they failed to do so after NOTCH2 depletion in the cancer cells (Figure 3D, Figure S3C-E). Moreover, knocking down JAG1 in the THP-1-derived macrophages (Figure S3E) significantly attenuated the survival of OVCAR8 cancer cells to a level similar to that of NOTCH2 knockdown in the cancer cells (Figure 3D, Figure S3C). And concurrent knock down NOTCH2 in cancer cells and JAG1 in macrophages did not cause further growth inhibition (Figure 3D).

To examine the macrophage involvement in taxol resistance *in vivo*, we depleted macrophages in ID8 syngeneic mouse models of ovarian cancer using liposomes-Clodronate (Figure S3F). Similar to the ovarian cancer patient samples discussed in Figure 3A-C, JAG1 is mainly expressed on macrophages from the ascites of the ID8 mouse model (Figure S3G, H). In the absence of macrophages, the *in vivo* tumors were substantially more sensitive to paclitaxel-induced growth arrest, and knocking down NOTCH2 in the tumor cells did not further enhance the tumor sensitivity to paclitaxel (Figure 3E). Moreover, we observed that upon the *in vivo* depletion of macrophages, the percentages of tumor cells with NICD2-positive nucleus significantly decreased (Fig. 3F), further supporting that macrophages were crucially involved in activating NOTCH2 signaling when the tumor cells were treated with paclitaxel. Overall, our data from both the *in vitro* and *in vivo* analysis suggest that tumor-macrophage interaction is the primary activator of NOTCH2-JAG1 juxtacrine signaling in the TME that mediates tumor resistance to paclitaxel.

### NOTCH2 signaling activates pro-survival pathways and stimulates the recruitment of pro-tumor macrophages

To pinpoint how activation of NOTCH2 signaling confers taxol resistance at the molecular and cellular level, we investigated the downstream signaling pathways of NOTCH2 using transcriptome analysis of NOTCH2-depleted cells. Principal component analysis of the transcriptome data from NOTCH2-knockdown OVCAR8 cells using two different NOTCH2 shRNA constructs gave consistent results (Figure 4A, Figure S4A). In addition to downregulation of the pro-survival PI3K-AKT signaling and MAPK signaling in the NOTCH2-knockdown cells, which indicated their involvement in NOTCH2-mediated paclitaxel resistance (Figure S4B), we found considerable downregulation of cytokines, such as CSF1 and IL-1β, upon NOTCH2 knockdown (Figure 4B). Further GSEA pathway analysis revealed that cytokine-cytokine receptor interaction pathway was indeed significantly downregulated under NOTCH2 knockdown (Figure 4C). We subsequently confirmed, by qPCR (Figure 4D, Figure S4C, D) and ELISA (Figure 4E), that taxol treatment induced expression of CSF1 and IL-1β, and NOTCH2 depletion markedly reduced their expression in both OVCAR8 and A549 cells. Moreover, overexpression of NICD2 rescued the expression of CSF1 and IL-1β under NOTCH2 depletion (Figure 4D). Analysis of the TCGA dataset also showed strong correlation of NOTCH2 level with CSF1/IL-1β levels (Figure S4E) in ovarian and lung cancers.

**Figure 4.**
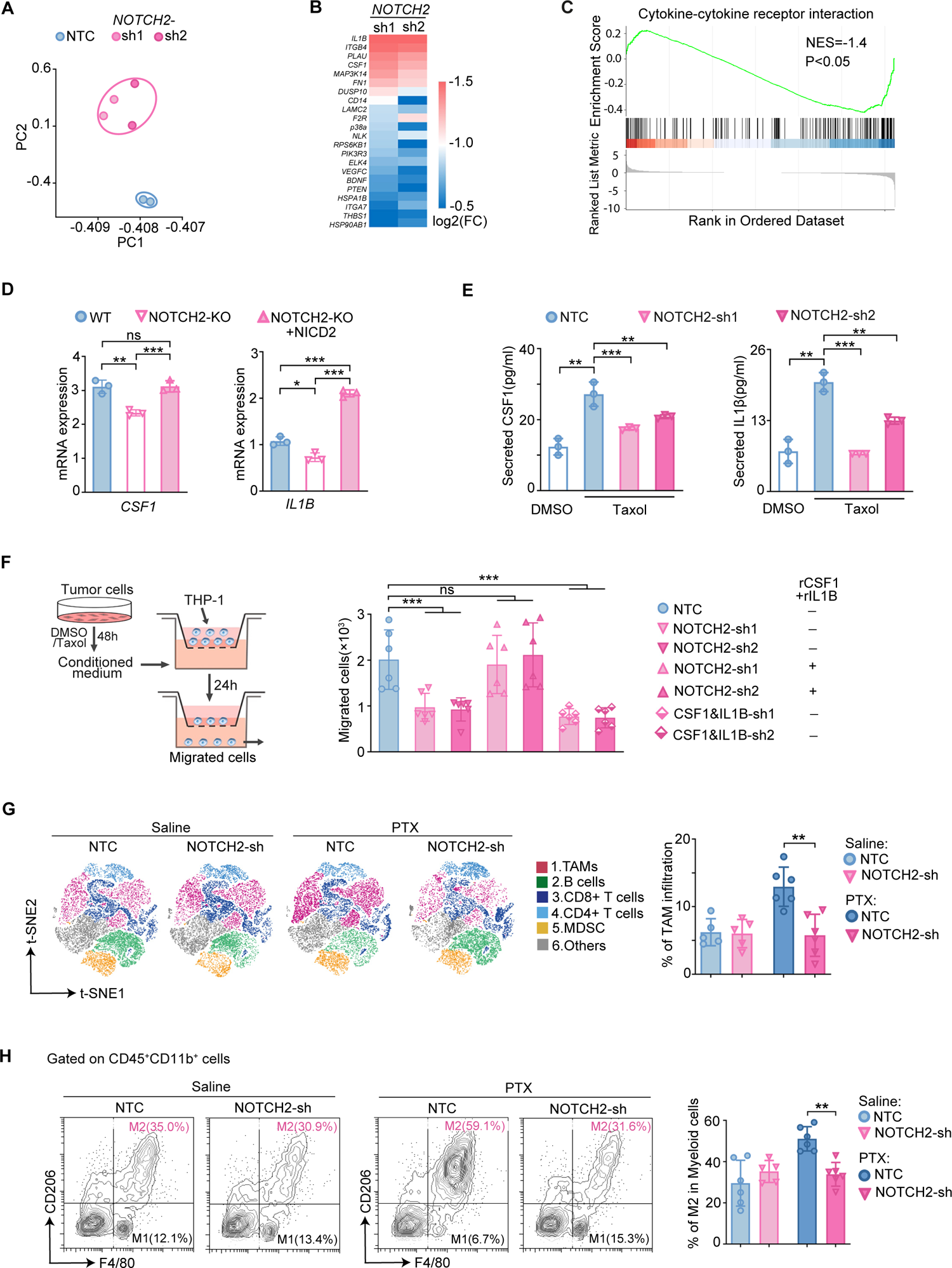
NOTCH2 upregulation activates pro-survival pathways and stimulates the recruitment of pro-tumor macrophages. **(A)** Principal-component analysis (PCA) of gene expression for wildtype and NOTCH2-knockdown OVCAR8 cells. Two different NOTCH2 shRNA clones were used. **(B)** Heatmap showing log2 (Fold-change) value (≥0.5) of DEGs enrichment of different signaling pathways. **(C)** Gene Set Enrichment Analysis (GSEA) of the RNA-Seq data. **(D)** RT–PCR analysis of *CSF1* and *IL1B* mRNA levels in A549 cells of NOTCH2-WT or -KO transfected with vector or NICD2 expression construct. Cells were treated with 100 nM taxol for 48 hrs. **(E)** ELISA-based quantification of the secreted CSF1 and IL1β in the medium of OVCAR8 cells with or without NOTCH2 knockdown. Cells were treated with DMSO or 10 nM taxol for 48 hrs. **(F)** Chemotactic with either vehicle or 10 nM taxol for 48 hrs were evaluated by Boyden chamber assay. Diagram of the experimental setup is shown on the left, and the quantified number of migrated cells is shown on the right. **(G)** t-SNE plot showing a low-dimensional representation of the different immune cell types in the ascites of ID8-carrying mice after one week of treatment. **(H)** FACS analysis of the ratios of M2 macrophages among the myeloid cells (CD45+CD11b+) from the ID8 mouse ascites. Data are shown as mean ±SD. Data of (D), (E) and (F) were obtained from three experimental replicates. p values were determined by two-tailed Student’s t test (ns, not significant; *p < 0.05, **p < 0.01, ***p < 0.001).

CSF1 and IL-1β are known to recruit pro-tumor M2 macrophages [19, 34], implicating a possible immunomodulatory role of NOTCH2 activation to increase pro-tumor macrophages and augment the pro-survival tumor-macrophage interactions in the TME. To characterize the effect of NOTCH2 activation in recruiting macrophages, conditioned medium from control and NOTCH2-depleted OVCAR8 cells were collected and their effects on the migration of THP-1 were measured by a transwell assay (Figure 4F). Compared to medium from the taxol-treated wildtype cells, medium from the NOTCH2-depleted cells under taxol treatment showed significantly less effect in promoting THP-1 migration (Figure 4F). There was no such difference for medium from DMSO-treated wildtype and NOTCH2-depleted cells (Figure S4F). Moreover, addition of recombinant CSF1 and IL-1β to the medium of NOTCH2-depleted cells resulted in an extent of THP-1 migration similar to that induced by the wildtype medium (Figure 4F). And concurrent depletion of CSF1 and IL-1β in cancer cells significantly reduced the migration of THP-1 cells (Figure 4F and S4G). Together these data illustrate that NOTCH2 signaling stimulates macrophage recruitment via upregulating cytokines, such as CSF1 and IL-1β.

We next examined the role of NOTCH2 in macrophage recruitment in ID8 syngeneic mouse model of ovarian cancer under paclitaxel treatment (Figure 4G, H and Figure S4I). Depletion of NOTCH2 by shRNA did not affect tumor growth in the control mice, but significantly attenuated the tumor growth in the taxol-treated mice (Figure S4I), confirming that NOTCH2 is involved in promoting taxol resistance *in vivo*. We noted that only the number of macrophages, but not the other immune cell types, was significantly reduced in mice bearing NOTCH2-depleted tumor cells upon taxol treatment (Figure 4G, H). Further analysis showed that these macrophages are mainly of the M2-type (Figure 4H). These *in vivo* data indicate that the NOTCH2-mediated paclitaxel resistance is enhanced by the macrophage recruitment.

### NOTCH2 inhibition sensitizes tumor response to paclitaxel in multiple mouse tumor models

Given that NOTCH2 upregulation and tumor-macrophage interaction via NOTCH2-JAG1 juxtacrine signaling drive the tumor resistance to paclitaxel, we next explored the combinatorial effect of NOTCH2 inhibition and paclitaxel in mouse tumor models. Mouse xenograft tumor model derived from OVCAR8 cells (Figure 5A) and A549 cells (Figure S5A) showed that NOTCH2 knockdown significantly attenuating tumor growth under paclitaxel treatment. Notably, knocking down NOTCH2 significantly reduced the number of tumor-infiltrated macrophages under taxol treatment (Figure S5B), confirming the *in vivo* role of NOTCH2 in promoting macrophage recruitment to the TME. As there is no NOTCH2-specific small-molecule inhibitor currently available, we used a pan-NOTCH inhibitor, RO4929097 (a Gamma-secretase inhibitor (GSI)), as a proof-of-concept. NOTCH2 inhibition by RO4929097 treatment was confirmed by the reduced expression of the NOTCH2 target gene, HES1 (Figure S5C). In the mouse xenograft tumor models of OVCAR8 (Figure 5B) and A549 (Figure S5D), combinatorial treatment of RO4929097 plus paclitaxel significantly enhanced tumor growth inhibition, as compared to paclitaxel or RO4929097 treatment alone. This synergistic inhibitory effect was not further enhanced when NOTCH2 was depleted, indicating that NOTCH2 is primarily responsible for the pan-NOTCH inhibition effect of RO4929097 (Figure S5E). We also evaluated the combinatorial treatment effect in the ID8 syngeneic mouse model (Figure 5C). Compared to paclitaxel treatment alone, RO4929097 plus paclitaxel showed significantly higher attenuation of tumor growth (Figure 5C) as well as reduction of macrophages in the tumor (Figure 5D), illustrating that blocking NOTCH2 signaling damped the recruitment of pro-tumor macrophages under taxol treatment.

**Figure 5.**
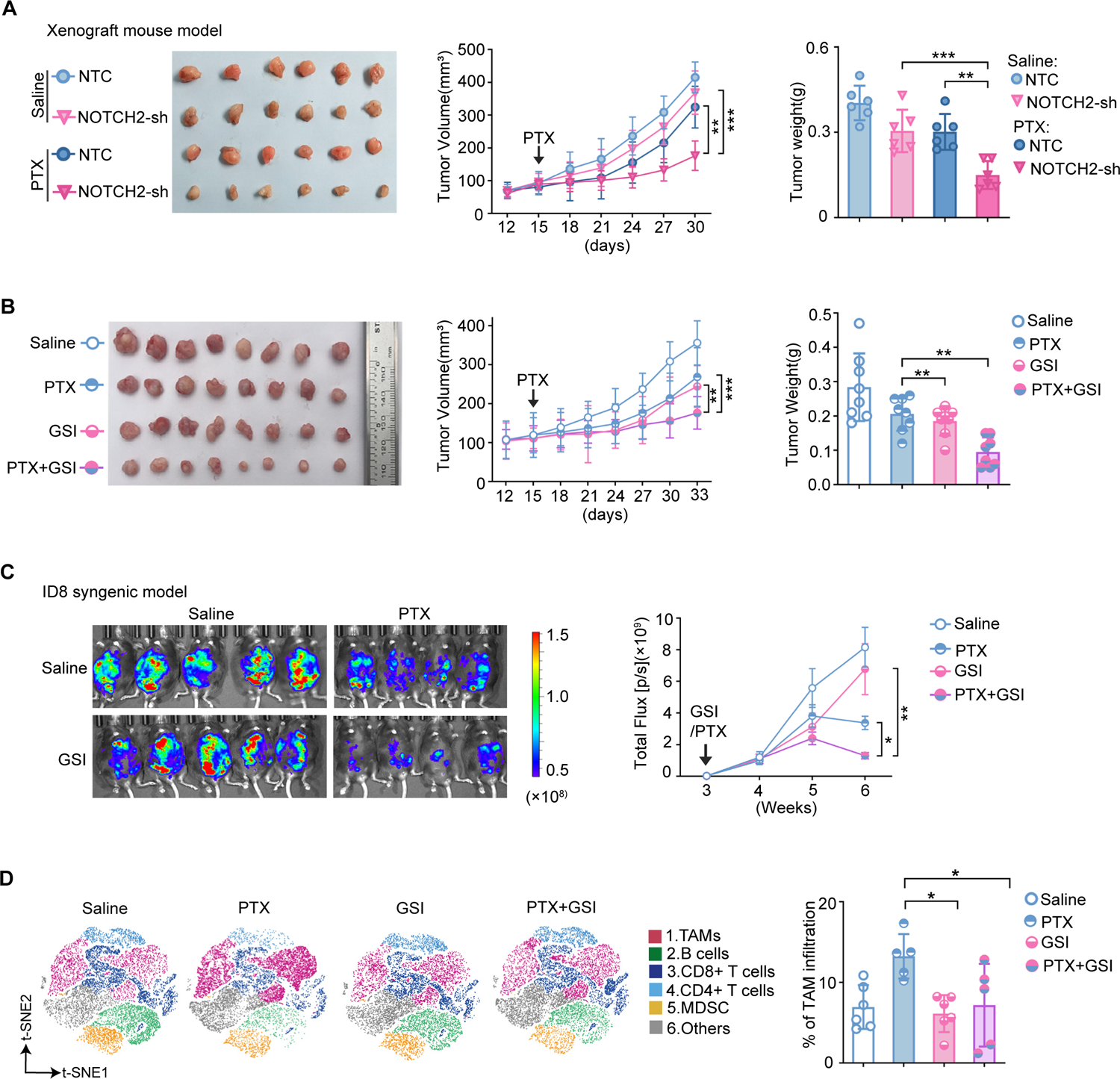
NOTCH2 inhibition sensitizes tumor response to paclitaxel in multiple mouse tumor models. **(A)** Representative dissected tumors (left), tumor growth curve (middle) and tumor weight (right panel) of OVCAR8-xenograft mice. OVCAR8 cells with doxycycline inducible NOTCH2 knockdown (sh) or non-target control (NTC) were injected and 10 mg/kg paclitaxel was administered by intraperitoneal injection when tumor volume reached about 100 mm^3^. Error bars are SEM. **(B)** Representative dissected tumors (left), tumor growth curve (middle) and tumor weight (right panel) from mice carrying OVCAR8 xenografts that were treated with GSI and paclitaxel. **(C)** Representative images of mice carrying ID8-luciferase cells three weeks after treatment (left panel) and the corresponding tumor growth curves (right panel). **(D)** t-SNE plot showing a low-dimensional representation of the different immune cell types in the ascites of ID8-carrying mice after one week of treatment. p values were determined by two-tailed Student’s t test (*p < 0.05, **p < 0.01, ***p < 0.001).

### NOTCH2 upregulation is associated with chemoresistance in ovarian cancer patients

To evaluate the clinical relevance of NOTCH2, we examined its level in tumors from patients treated with taxanes. Taxol-based chemotherapy is frequently used for advanced ovarian cancer, and we analyzed paired primary tissues and recurrent surgical tissues from ovarian cancer patients by immunochemical staining (Figure 6A). The IHC score (Figure S6A, defined by expression levels as detailed in methods) revealed a significant correlation between NOTCH2 level and tumor recurrence in these patients after paclitaxel/platinum treatment (Figure 6A, B). For patients that relapsed after adjuvant therapy, progression-free survival rate was substantially lower for those with tumors showing high expression of NOTCH2 (Figure 6C). Such correlation was not observed for patients without taxol-based adjuvant therapy, suggesting that NOTCH2 expression is particularly associated with tumor recurrence after taxol-based treatment (Figure 6C). In addition, analysis of ovarian cancer patient data from the TCGA database confirmed that the overall survival rate is low for high NOTCH2 expression, but not for the other NOTCH receptors, in the taxol treatment cohorts (Figure S6B). Together these clinical data from patients indicate that NOTCH2 levels are significantly correlated with the tumor sensitivity to taxol treatment for ovarian cancer patients. NOTCH2 could be employed as a viable therapeutic target and prognostic biomarker for predicting patient responses to taxane-based therapy.

**Figure 6.**
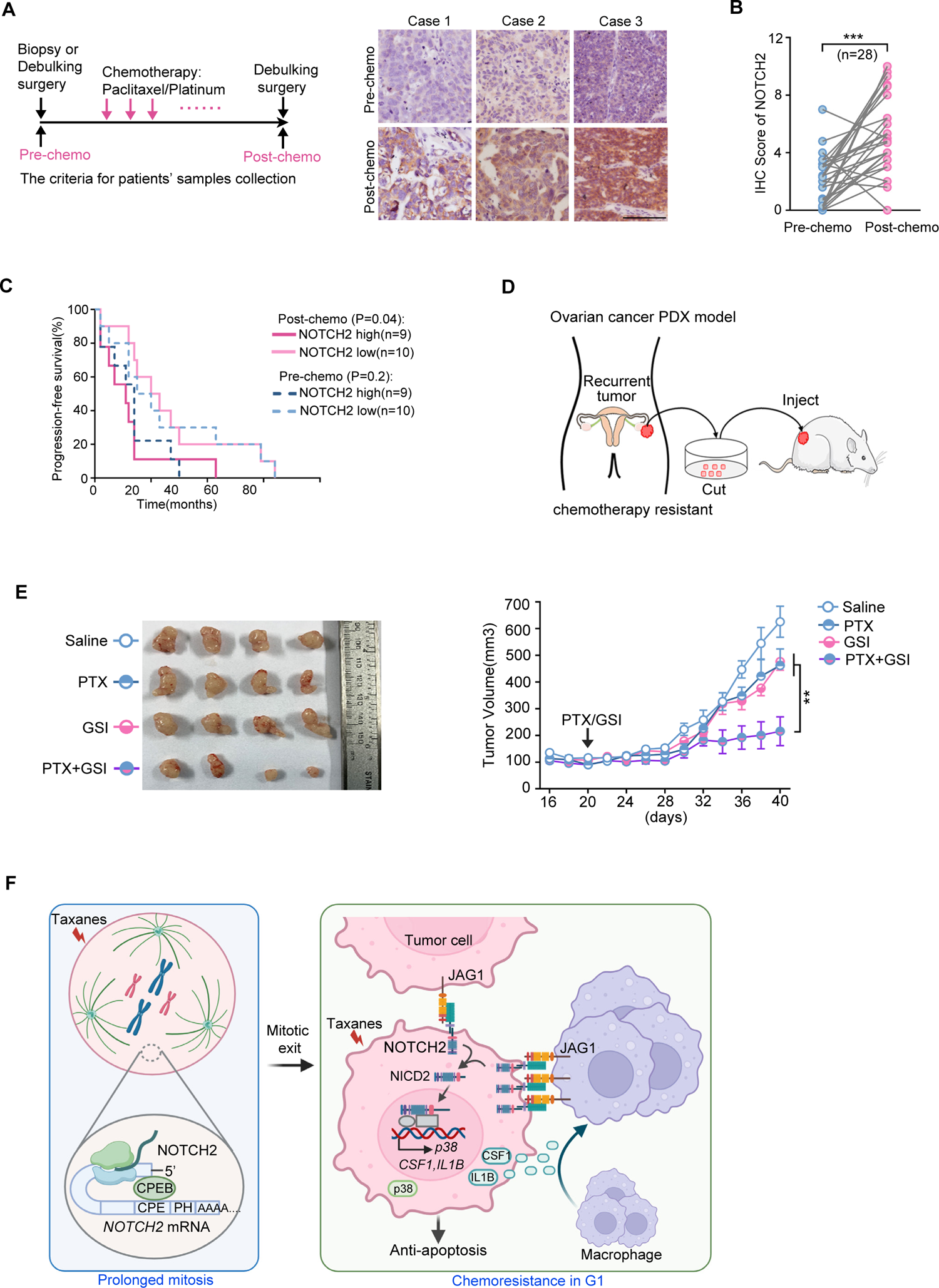
High NOTCH2 level is associated with chemoresistance of clinical samples. **(A)** The criteria for the collection of patient samples and representative NOTCH2 staining in matching images of tumor specimens from ovarian cancer patients obtained pre (chemotherapy)- and post (chemotherapy)-taxane therapy. Scale bars are 200 μm. **(B)** The IHC score of NOTCH2 was quantified. **(C)** Kaplan-Meier survival analysis of ovarian cancer patients. Patients were divided according to the IHC score of NOTCH2 (cutoff: median) in tumor tissues of pre (chemotherapy)- and post (chemotherapy)-taxane therapy, respectively. **(D)** Schematic diagram depicting the ovarian cancer PDX model derivation. **(E)** Effect of GSI, paclitaxel and their combination on tumor growth of ovarian cancer PDX mice. Representative tumors (left panel) and tumor volume (right panel) for each group are shown. **(F)** Proposed working model. Our results revealed that NOTCH2 translation was upregulated during prolonged mitosis and the NOTCH2-JAG1 juxtacrine signaling between tumor and macrophages was strongly activated in the post-mitotic G1 phase, which confers taxol resistance. Error bars are SEM. p values were determined by two-tailed Student’s t test (**p < 0.01, ***p < 0.001).

To further test the efficacy of the combinatorial treatment of NOTCH inhibitor and paclitaxel in a clinically relevant model, a patient-derived xenograft (PDX) model for ovarian cancer was generated (Figure 6D). The tumor tissues were from a patient with recurrent tumor after adjuvant chemotherapy with paclitaxel/platinum (Figure S6C), and the PDX was validated by staining multiple ovarian cancer markers (Figure S6D). Upon combinatorial treatment of paclitaxel and RO4929097, robust synergistic effect in suppressing tumor growth was observed in the PDX tumor (Figure 6E). The *in vivo* data from multiple mouse tumor models all point to NOTCH2 inhibitor as a promising combinatorial therapy to reverses paclitaxel resistance in NOTCH2-positive tumors.

## Discussion

Chemoresistance is a major challenge for cancer treatment and mechanistic understanding of the molecular and cellular determinants underlying drug resistance is key to develop new combinatorial strategies to improve the clinical efficacy of chemotherapeutics for a broad patient population. By investigating contact-dependent interactions between cancer cells and macrophages in the TME that contribute to tumor resistance to paclitaxel, we identified a novel NOTCH2-JAG1 juxtacrine signaling in the TME that is druggable to sensitize tumor response to paclitaxel. The newly proposed therapeutic approach of NOTCH2 inhibition is possibly more tumor selective and can effectively disrupt pro-survival tumor-macrophage interaction without broadly affecting the normal macrophage functions at the system level. Moreover, we found NOTCH2 inhibition can reduce pro-survival signaling in tumor cells as well as attenuate the immuno-suppressive TME, i.e., by inhibiting the tumor infiltration of M2-type macrophages. Such effect on blocking both intrinsic and extrinsic pro-tumor factors makes NOTCH2 an attractive combinatorial target to reverse paclitaxel resistance.

On the mechanism of paclitaxel resistance (Figure 6F), we found the membrane receptor NOTCH2 is synthesized and upregulated during prolonged mitotic arrest induced by paclitaxel. Shortly after the cells exit mitosis and enter the next G1 phase, NOTCH2 is activated by its cognate activating ligand, JAG1, which was mainly expressed on the neighboring macrophages. Cleaved intracellular NOTCH2 then translocates to the nucleus and activates the transcription of a series of target genes. The downstream network activated by NOTCH2 includes both intracellular signaling that upregulates pro-survival pathway activities and extracellular signaling via cytokine production that reshapes the TME (Figure 6F). In particular, a positive feedback of tumor-macrophage interaction via the NOTCH2-JAG1 axis plays a key role in rendering taxol resistance. On the one hand, NOTCH2 activity increases the production of cytokines that recruit tumor-associated macrophages. The infiltrated macrophages, on the other hand, further enhances NOTCH2 activation via the JAG1 ligand highly expressed on their surface, therefore forming a positive feedback loop to promote NOTCH2-mediated pro-survival signaling (Figure 6F). As the transcription of NOTCH2 only starts a few hours after cells exit the prolonged mitosis (Figure S1C), the advantage of NOTCH2 upregulation at the translational level during mitotic arrest could be that it renders a much faster response from the pro-survival pathways, which occurs as soon as the cells exit the prolonged mitosis and form contacts with the neighboring cells that express JAG1. The rapid activation of NOTCH2-mediated pro-survival signaling may protect cancer cells from apoptosis that is initiated during the prolonged mitotic arrest. Overall, our results revealed that translational upregulation of NOTCH2 not only promotes cell survival via cell autonomous signaling but also triggers a non-cell autonomous positive feedback loop to recruit TAMs.

A number of pan-NOTCH and NOTCH ligand inhibitors have been tested either alone or in combination with taxanes in clinical trials for different cancer types [35–39]. The NOTCH inhibitor, Nirogacestat, received FDA approval in 2023 for treating Desmoid Tumors [40]. As our results showed NOTCH2 inhibition with paclitaxel showed synergistic effects (Figure 6), the combinatorial drug action of NOTCH inhibitor in attenuating tumor growth is likely mitosis dependent. In addition to non-specific GSIs for pan-NOTCH inhibition, monoclonal antibodies against individual NOTCH receptors are also currently being developed [32, 41]. Tarextumab, a human monoclonal antibody that inhibits both NOTCH2 and NOTCH3, suppressed tumor growth in breast and ovarian tumor models when used in combination with taxol [42]. However, this combination failed in clinical trials for pancreatic cancer [43]. This might be due to the side effects that limited the viable clinical dose, and our results suggest that a NOTCH2-specific inhibitor is likely more effective for combinatorial therapy with anti-mitotic drugs. To develop more specific therapeutics that selectively target NOTCH2, biologic antibody against NOTCH2 may render higher efficacy than small-molecule inhibitors and dual-targeted antibodies to enhance taxane response for patients with NOTCH2-positive tumors.

## Methods

### Cell lines and cell culture

All cell lines were obtained from the sources indicated in the key resources table (Table S1). HeLa, A549, OVCAR8, ID8, HEK293T and HeyA8 were cultured in DMEM (Invitrogen/Thermo Fisher Scientific, MA, USA) supplemented with 10% fetal bovine serum (FBS) (Gibco, Grand Island, NY, USA) and 1% penicillin/streptomycin (P/S) (Beyotime Biotechnology, Jiangsu, China) at 37°C with 5% CO2. THP-1 cells were cultured in RPMI-1640 (Invitrogen/Thermo Fisher Scientific, MA, USA) supplemented with 10% FBS and 1% P/S. Cells were tested regularly for mycoplasma.

### Live cell imaging

For live cell imaging, 2×10^4^ cells were seeded into 8 well-Sliders (#80826, ibidi) and maintained at 37°C overnight in DMEM with 10% FBS and 1% P/S. The medium was changed to phenol red-free L-15 medium (Invitrogen/Thermo Fisher Scientific, MA, USA) with 10% FBS and 1% P/S. Drugs or DMSO were added before the start of the imaging experiments. Time-lapse images were acquired at 5 min intervals with a 20×objective lens mounted on an Eclipse Ti microscope (Nikon, Tokyo, Japan). Time of mitosis and interphase was determined morphologically (round-up morphology is defined as mitosis and flat morphology is defined as interphase). The onset of cell death was scored by the characteristic morphological changes of blebbing and cell lysis.

### Cell synchronization

HeLa, A549 and OVCAR8 cells were synchronized by double thymidine block (2 mM, Sigma). Specifically, cells were plated in 10 cm dishes one day before treatment. Then 2 mM thymidine was added to the dishes to arrest the cells for 19 hours (hrs). After washing the dishes three times with PBS, cells were released into fresh culture medium for 9 hrs, followed by a second thymidine block for 16 hrs. Then the dishes were washed three times with PBS to release the cells to complete DMEM again. 9 hrs later, DMSO and 5 nM paclitaxel were added to each group, respectively. The mitotic cells were collected by shake-off. The mitotic cells from DMSO and taxol treatment were defined as mitosis (M) and prolonged mitosis (pM) cells, respectively.

### Ribosome profiling and mRNA sample preparation

For ribosome profiling, HeLa cells were synchronized by double thymidine block (2 mM, Sigma). Mitotic (M) and prolonged mitotic (pM) HeLa cells were lysed in lysis buffer (20 mM Tris pH 7.5, 150 mM KCl, 5 mM MgCl2, 1 mM dithiothreitol, 8% glycerol) supplemented with 0.5% Triton X-100, 30 U/ml Turbo DNase (Ambion, Life Technologies, CA, United States) and 100 μg/ml cycloheximide (Sigma Aldrich, MO, United States). Ribosome-protected fragments were then isolated and sequenced as previously described [44]. The resulting mRNAs were modestly fragmented by partial hydrolysis in bicarbonate buffer so that the average size of fragments was ∼80 bp. The fragmented mRNAs were separated by denaturing PAGE and fragments of 26–34 nt were selected. The libraries were prepared and sequenced as previously described [44].

### Western blot

Cells were seeded to cell culture dishes 24 hrs prior to treatment. To measure the apoptotic markers (Caspase-3 and PARP), the culture media were centrifuged at 1200 rpm for 5 mins to collect the apoptotic cells. The cells that remained attached to the dish were washed using PBS three times before adding lysis buffer. Both apoptotic and surviving cells were lysed using cell lysis buffer (50mM Tris HCl, pH 7.4, 250mM NaCl, 1 mM EDTA, 50mM NaF, 0.5% Triton X-100 and protease inhibitors), and then mixed together. For other experiments, the culture media were aspirated and the cells in the dishes were washed with PBS three times before adding lysis buffer. Cells were lysed for 30 mins on ice before centrifugation at 12000 rpm for 15 mins at 4°C. The supernatants of the samples were collected. The protein concentration of each sample was determined using Bradford Kit (Sangon Biotech. Shanghai, China). For immunoblotting, proteins were resolved by SDS-PAGE and transferred onto polyvinylidene difluoride (PVDF) membranes (Millipore). Immunoblots were developed using Western Lightning Chemiluminescence Reagent Plus (Advansta, Menlo Park, CA, USA).

### Gene knockdown by shRNA

Lentiviral particles were packaged in HEK293T cells. The virus was harvested 48 hrs after transfection and filtered through non-pyrogenic filters with a pore size of 0.45 mm (Merck Millipore, Billerica, MA, USA). For infection, cells were seeded at 1-3 x10^5^ cells per well in a 35 mm dish. 24 hrs later, diluted lentivirus and 8 μg /ml polybrene (Sigma-Aldrich, St.Louis, MO, USA) were added to the cells. 24 hrs after infection (48 hrs for THP-1), the medium with viral particles was removed. For stable cell selection, puromycin (1 μg/ml) was added 48 hrs post infection.

### CRISPR-Cas9 gene targeting

One pair of guides against the exon-30 sequence of Human NOTCH2 was designed using the guide design tool found at: (https://zlab.bio/guide-design-resources/) with the input of Human NOTCH2 exon-30 sequence. Guide sequences were cloned into the pSpCas9 (BB)-2A-Puro vector that was previously described [45]. Lipo3000 was used to transfect the cells, and puromycin was used to select the transfected cells. Cells were selected for five days after transfection, then plated for single colonies. Multiple candidate clones were picked and tested for gene disruption by western blot. The ones in which NOTCH2 protein expression was downregulated were selected for genomic extraction and PCR. The PCR fragment was sent to SANGON for sequencing to confirm that NOTCH2 exon-30 was disrupted. The CRISPR-guide sequences and vector information were listed in the Key Resources Table.

### RNA Immunoprecipitation (RIP)

RIP was carried out as previously described [46]. Briefly, HeLa cells in prolonged mitosis were crosslinked in a UV cross-linker (UVP) at 120 mJ cm^-2^ strength for 2 mins. Cells were then harvested in ice-cold RIPA buffer (50 mM Tris-Hcl, pH 8.0), 150 mM NaCl, 5 mM EDTA, 1% NP-40 and 1% SDS) and sonicated for 5 mins with a Sonics Vibra-Cell with a 3-second on and 6-second off procedure. The cell suspension was centrifuged at 13,000×g for 20 mins at 4°C, and the supernatant was collected. Antibody or IgG (as control) was added and incubated for 4 hrs at 4°C with Protein G Dynabeads (Life Technologies) for antigen coupling. Suspension of sample was added next and allowed to bind for at least 4 hrs at 4°C. The protein antibody-bead complexes were washed twice with RIPA buffer and twice with RIPA buffer supplemented with 500 mM NaCl. One-fifth of the beads after the last wash were heated at 100°C for 10 mins in SDS-loading buffer and then saved for western blotting. The remaining protein antibody-bead complexes were digested with proteinase K at 55°C for 30 mins, followed by extraction with Trizol Reagent (Takara, Dalian, China) to obtain RNA.

### RNA-Seq and KEEG pathway enrichment analysis

RNA was extracted from OVCAR8 cells using Trizol Reagent (Invitrogen, Carlsbad, CA, USA) according to the manufacturer’s protocol. Subsequently, total RNA was quantified using a Nano Drop and Agilent 2100 bioanalyzer (Thermo Fisher Scientific, MA, USA). The Beijing Genomics Institute (BGI, Shenzhen, China) performed the RNA integrity checking, library preparation and sequencing. Oligo (dT)-attached magnetic beads were used to purify mRNA. The purified mRNA was fragmented into small pieces with fragment buffer at appropriate temperature. Then cDNA was generated according to the manufacturer’s protocol by BGI. The cDNA product was validated on the Agilent Technologies 2100 bioanalyzer for quality control. The final library was amplified with phi29 to make DNA nanoball (DNB) which had more than 300 copies of one molecule. DNBs were loaded into the patterned nanoarray and single-end 50 bases reads were generated on BGIseq500 platform (BGI-Shenzhen, China).

Sequence reads were aligned to the human reference genome with GRCh38.p11 (hg38). Differential Expression Genes (DEGs) were analyzed using the DEseq2 R package. KEEG pathway Enrichment Analysis was performed using KOBAS (http://kobas.cbi.pku.edu.cn/kobas3/genelist/). The input gene lists were generated from the overlapping of differentially downregulated genes in two comparisons based on adjusted p value < 0.05 and log2 Fold Change < −0.5 (NOTCH2-sh1 compared to NTC, NOTCH2-sh2 compared to NTC), which consists of 522 significantly downregulated genes in total. Cut-off values of FDR < 0.05 were used to select the top enriched pathways. Original data are available in the NCBI Gene Expression Omnibus (GEO) under the accession number GSE158569.

### Gene expression analysis by qRT-PCR

qRT-PCR was conducted as previously described [47]. Briefly, total RNA was extracted from cells using Trizol Reagent (Invitrogen, Carlsbad, CA, USA) following the manufacturer’s protocol. Extracted RNA samples were quantified using Nanodrop (Life Technologies, Carlsbad, CA, USA). cDNA was prepared from 1 μg RNA using Oligo (dT) and Superscript III reverse transcriptase according to the manufacturer’s protocol (Vazyme, China). Quantitative PCR was performed using SYBR^®^ Premix (Vazyme, China). Data were collected using a PIKOREAL 96 real-time PCR system (Thermo Scientific, MA, USA). All reactions were performed in triplicate.

The primers used in this study are listed in the key resources table (Table S1). The primers were designed based on PrimerBank or OriGene (OriGene Technologies, Inc.) and were blasted to confirm the specificity using NCBI primer blast. The primers were synthesized by SANGON (Shanghai, China).

### Soluble JAG1 ligand immobilization and cell stimulation

The recombinant rat JAG1-Fc fusion chimera, goat anti-Human lgG FC antibody and recombinant human lgG1-FC (R&D Systems, Minneapolis, MN) were dissolved in PBS at a final concentration of 10 μg/mL. Soluble JAG1 ligand and the control for JAG1-human IgG-Fc immobilization was performed according to previous study [48, 49]. Briefly 2 μg or 0.5 μg goat anti-Human lgG FC antibody were added to 12-well plate or 8 well-Sliders per well and incubate for 2 hrs at room temperature. The solution was then aspirated and washed with PBS three times. Then 1 μg or 0.25 μg recombinant rat JAG1-Fc fusion chimera and recombinant human lgG1-FC were added to the coated 12-well plate or 8 well-Sliders, respectively, and incubated at 4°C for 20 hrs. The solution was aspirated and washed with PBS before plating at 1×10^5^ cells or 1×10^4^ cells per well.

### ELISA

Cells were seeded in a 60 mm dish with 3 ml complete culture medium 24 hrs prior to treatment. To determine IL-1β and CSF1 levels in the cell culture medium, following the indicated treatments for 72 hrs, conditioned media were centrifuged at 1000 ×g for 10 mins and supernatants were collected to determine the cytokine levels. Cytokine levels were measured using ELISA kits (ABclonal) according to the manufacturer’s instructions. Cell numbers were counted to normalize the total level of the cytokines.

### THP-1 differentiation and co-culture assays

M2-polarized THP-1 macrophages were generated as described previously [50]. Briefly, 2×10^6^ THP-1 cells were seeded in a 35 mm dish and treated with 320 nM PMA for 6 hrs, then cultured with PMA plus 20 ng/ml IL-4 and 20 ng/ml IL-13 for another 18 hrs. For the co-culture assay, OVCAR8-GFP stable cells were seeded with M2 macrophages (ratio 1:1) in 6-well plates 24 hrs prior to adding taxol. The density of the OVCAR8-GFP cells was captured using an Olympus microscope with a GFP channel every day for 3 days. The confluence of OVCAR8-GFP cells was quantified using image J (Analyze>Analyze particles).

### *In vivo* mouse tumor models

All animals were maintained in the house under specific pathogen-free conditions with unrestricted access to food and water for the duration of the experiments. All mouse experiments were approved by The Ethics Committee of the University of Science and Technology of China (2019-N(A)-059). Balb/c mice (athymic nu/nu, 6-8 weeks old, SLAC Laboratory Animal) were subcutaneously injected with 5×10^5^ OVCAR8 or A549 cells. For NOTCH2 knockdown, OVCAR8 cells that stably express inducible NOTCH2-shRNA were injected and 1mg/ml doxycycline (Dox) in the drinking water was fed to mice during the experiments. For experiments with NOTCH inhibitor, the mice were randomly divided into different groups when the size of the tumor reached 100-150 mm^3^. Paclitaxel (10 mg/kg) were administered by intraperitoneal (i.p.) injection every two days. RO4929097 (3 mg/kg) were administered orally every two days. Tumor growth was recorded by blind measurement of two perpendicular diameters of the tumors and tumor size was calculated using the formula 4p/3 × (width/2)^2^ × (length/2). Tumors were harvested after treated for two weeks (experimental endpoint).

For the murine ID8 model, 8-10 week-old female C57BL/6 mice (SLAC Laboratory Animal) were injected intraperitoneal (i.p) with 5×10^6^ ID8 cells. Six weeks after tumor inoculation, ascitic fluid was collected for immunostaining.

To generate the patient-derived xenograft (PDX) model, fresh tissues from patients with histologically confirmed serous ovarian cancer were collected at the time of debulking surgery at the first affiliated hospital of University of Science and Technology of China (USTC). All surgically resected tumors were collected with written patient consent and following the institutional review board approved protocols of USTC. Patient heterotransplant number was used to protect patient identity. Nonnecrotic adjacent malignant tissue was procured by clinical staff with specialized training in gross dissection. The tumor tissue was minced on ice. Approximately 0.3 to 0.5 cm^3^ of tumor slurry was mixed with 1:1 Matrigel (Corning) before being injected into at least three female NOD-SCID mice subcutaneously. Moribund mice with tumors were sacrificed, and to maintain each tumor graft model as the low passage, tumor from the initial founder mouse/mice was expanded with a single passage into 10+ mice to generate sufficient tumor volume for banking and future experiments. If the founder tumorgraft volume was insufficient, a pre-expansion amplification of tumor volume may be necessary for a small cohort of mice. Cryopreserved tumors were minced and stored at 1:1 ratio in freezing media (90% FBS and 10% dimethyl sulfoxide) overnight at −80°C and then in liquid nitrogen indefinitely. Macrophages in C57BL/6 mice were depleted using clodronate-containing liposomes according to a published protocol [51]. Mice were intravenously injected one dose of clodronate-liposomes (0.2 ml/mouse at 5 mg/ml, From Vrije Universiteit Amsterdam).

### Flow cytometry analysis (FACS)

Ascites were collected and incubated in Red Blood Cell Lysis Buffer (Beyotime Biotechnology) for 10-15 mins at 4°C to lyse the red blood cells. Cells were washed and counted using automated cell counter (Countstar). 1×10^6^ cells were blocked with FcR Blocking Reagent (BioLegend) or mouse serum, and stained for cell surface markers. All antibodies were purchased from BioLegend. Flow cytometry acquisition was performed on CytoFLEX (BD Biosciences), and data were analyzed using CytExpert software (2.4.0.28). Changes in the proportion of tumor-infiltrating immune cell subsets were compared by dimensionality reduction using t-SNE analysis using R (version 4.3.0).

### Chemotaxis assay

For analyzing cell migration, THP-1 cells were seeded (10^5^ cells/ 100 µl DMEM containing 0.1% FBS) onto the top chamber of transwell filters (3-µm; Corning). Filters were placed in a 24-well plate that contains conditioned medium isolated from the vehicle- or taxol-treated (5 nM) OVCAR8 cells. 24 hrs following the incubation, cells in the lower chamber were isolated and counted. Samples were run in triplicates for each group.

### Immunofluorescence

Immunofluorescence was conducted as previously described [52]. For cell lines, cells were plated onto 22×22 mm coverslips 24 hrs prior to treatment. Cells were washed with warmed PHEM (60mM PIPES (pH 6.8), 25mM HEPES, 10 mM EGTA and 2 mM MgCl2) and fixed in 4% formaldehyde for 15 mins at room temperature. For tumor tissues, the tissues were fixed in 4% formaldehyde for 14 hrs at 4°C prior to dehydrating by 30% sucrose for 24 hrs at 4°C. The tissues were then embedded into OTC gel and stored at −80°C before dissection for slides. For human or mouse ascites, fresh ascites were centrifuged at 2000 rpm for 5 mins at 4°C to collect cells and spread on Poly-L-Lysine coated slides. 10 mins later, when the ascites on slides were dried, cells were fixed in 4% formaldehyde for 15 mins at room temperature. Then cells or tissues were extracted by 0.2% Triton X-100 in PHEM, blocked with 1% bovine serum albumin in TBST for 30 mins, incubated with primary antibody for 2 hrs at room temperature, washed with TBST three times and incubated with secondary antibodies for an additional 1 hr at room temperature. DNA was stained with 4, 6-diamidino-2-phenylindole (DAPI) for 3 mins. Coverslips were mounted using ProLong antifade (Sigma). Images were acquired using a DeltaVision microscope (GE Healthcare, Buckinghamshire, UK) with a 60× or 20× objective lens, and optical sections were acquired 0.3-0.5 μm apart in the z-axis and shown as maximal intensity projections.

### Histology and Immunohistochemistry (IHC)

Tissue specimens from mice were fixed in 10% buffered formalin for 24 hrs and stored in 70% ethanol until paraffin embedding. 5 μm sections were stained with hematoxylin and eosin (HE) or used for immunohistochemical studies.

Human OV tissue samples were obtained from The First Affiliated Hospital of USTC. All samples were obtained with informed consent, and the study was approved by the Ethics Committee of USTC (2019-N(H)-212), as well as the principles expressed in the Declaration of Helsinki. Patients’ characteristics are shown in Table S2.

Immunohistochemistry was performed on formalin-fixed, paraffin-embedded mouse and human tissue sections using a biotin-avidin method as described before [47]. Sections were developed with DAB and counterstained with hematoxylin. Images were taken using a Nikon Ti microscope equipped with a 20× lens. Quantification of NOTCH2 IHC chromogen intensity was performed as previously described [53]. Briefly, staining intensity (0, 1, 2, 3) and percentage of positive cells among cancer (0-25% recorded as 1, 25-50% as 2, 50-75% as 3 and >75% as 4) were evaluated by a pathologist in USTC. The final immunoreactive scores were calculated by multiplying the two numbers as described before [53].

Multiplexed IHC (mIHC) assay to examine CD68 and JAG1 expression in human OV tissue samples was conducted according to the protocol of DendronFluor TSA kit (Jilin Histova Biotechnology company, China).

### TCGA and GEO data analysis

Clinical data and RNA expression data with normalized RSEM values of the serous ovarian cancer dataset (TCGA-OV) were retrieved from FireBrowse (http://firebrowse.org). 343 ovarian cancer patients had associated comprehensive follow-up records, RNA expression data and records of clinical treatment.

NOTCH receptors and its ligand expression levels were compared using RSEM values. For the survival curves, TCGA-OV patients were divided into two groups: patients that were treated with taxane-based therapy, including paclitaxel and docetaxel, and those that did not receive taxane-based treatment, but received other types of therapies. The curated clinical information was recorded as previous work [54]. Kaplan-Meier survival curves were drawn with the surminer R package (https://CRAN.R-project.org/package=survminer). Patients were stratified along with the best-cutoff value. GSE40484 [55] and GSE126080 [56] were downloaded from NCBI-GEO datasets (https://www.ncbi.nlm.nih.gov/geo/). Expression profiles of macrophages sorted from ovarian cancer ascites of mice and human were analyzed by microarray. Expression levels of NOTCH receptors and ligands were extracted and analyzed using R package and Graphpad.

TIMER2.0 (http://timer.cistrome.org/), an online tool, was used to analyze tumor-infiltrated macrophages in TCGA-OV patients [57]. The results obtained from the traditional method-CIBESORT-ABS were extracted.

## Statistical analysis

All the data analyses were performed with GraphPad Prism statistical software. p value was determined by two-way ANOVA for tumor growth, or Log-rank test for survival, or two-tailed t-tests for other analyses. A value of p < 0.05 was considered statistically significant.

## Data and code availability

Ribosome profiling and RNA sequencing data generated in this study are available at the NCBI-GEO database with the project accession ID GEO: GSE158569. Previously published expression datasets used in this study are available through GEO under GEO: GSE40484 and GSE126080. The Cancer Genome Atlas (TCGA) data were downloaded from Firebrowse (http://firebrowse.org/).

## Ethics approval statement and Patient Consent statement

All mouse experiments were approved by The Ethics Committee of the University of Science and Technology of China (2019-N (A)-059) The study using patient samples was approved by the Ethics Committee of USTC (2019-N (H)-212). We have obtained the patients consent for the studies.

## Acknowledgments

We thank all members in the Yang lab for discussion. This work is supported by: National Science Foundation of China 32370776, 92357301 to ZY, 31970670, 32170736 to JG, 32000492 to FY; National Key R&D Program of China 2022YFA1303100 to ZY. The Fundamental Research Funds for the Central Universities YD9100002011 to Z.Y.; Research Funds of Center for Advanced Interdisciplinary Science and Biomedicine of IHM of USTC (QYPY20220017 to Z.Y.); and the Hong Kong Research Grant Council (#RFS2021-2S01) to JS. JS also knowledges the funding support from the “Laboratory for Synthetic Chemistry and Chemical Biology” under the Health@InnoHK Program, ITC, HKSAR.

## Conflict of Interest

The authors declare that they have no conflict of interest.

---

**Figure S1.**
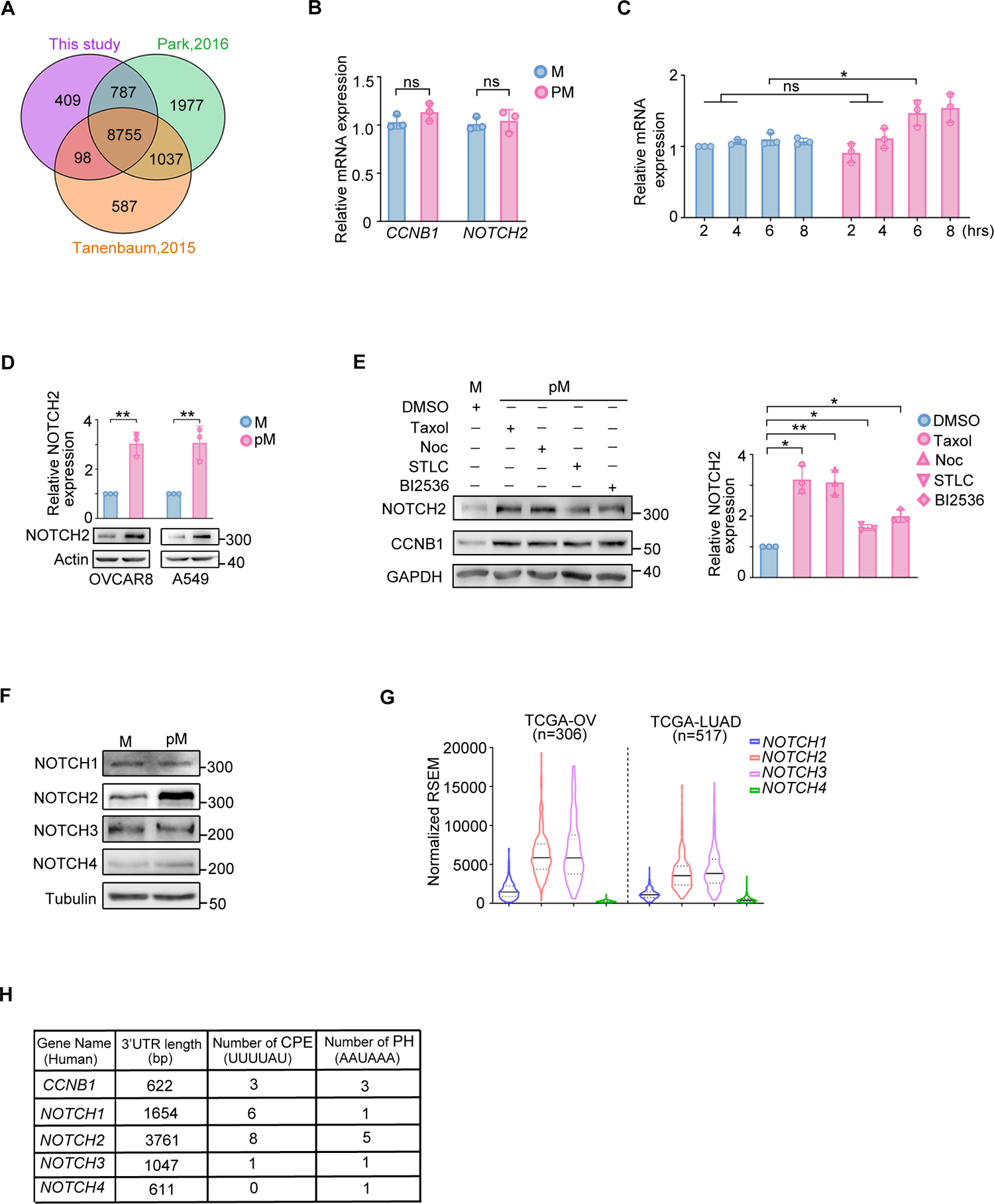
NOTCH2 translation is upregulated during taxol-induced prolonged mitosis. **(A)** Venn diagram showing the overlapping genes protected by ribosome during mitosis in this study, Park 2016, and Tanenbaum 2015. **(B)** RT–PCR detection of mRNA levels of the indicated genes in HeLa cells in mitosis (M) and prolonged mitosis (pM). **(C)** HeLa cells in mitosis (M) and prolonged mitosis (pM) induced by 0.5 μM Nocodazole were harvested by shake-off and re-plated in a culture dish with fresh medium for various hours before being collected. RT-PCR were conducted to analyze the mRNA levels of NOTCH2. **(D)** Western blot analysis of NOTCH2 in OVCAR8 and A549 cells in mitosis (M) and prolonged mitosis (pM). Quantified results are shown at the top. **(E)** Western blot analysis of NOTCH2 in HeLa cells in mitosis (M) and prolonged mitosis (pM). Cells in prolonged mitosis were induced by 5 nM taxol, 0.5 μM Nocodazole (Noc), 5 μM STLC and 100 nM BI2536, respectively. **(F)** Western blot analysis of NOTCH receptors in HeLa cells in mitosis (M) and prolonged mitosis (pM) cells. **(G)** Analysis of NOTCH receptors at the mRNA level in ovarian cancer and lung adenocarcinoma of NSCLC using TCGA data. **(H)** Characteristics of the NOTCH receptors and cyclin B1 transcripts’ 3’-UTR. The length and number of cytoplasmic polyadenylation elements (CPE) and polyadenylation hexanucleotide (PH) are presented. Data are shown as mean ±SD, and data of (B), (C), (D) and (E) were averaged from three experimental replicates. p values were determined by two-tailed Student’s t test (ns, not significant; *p < 0.05, **p < 0.01).

**Figure S2.**
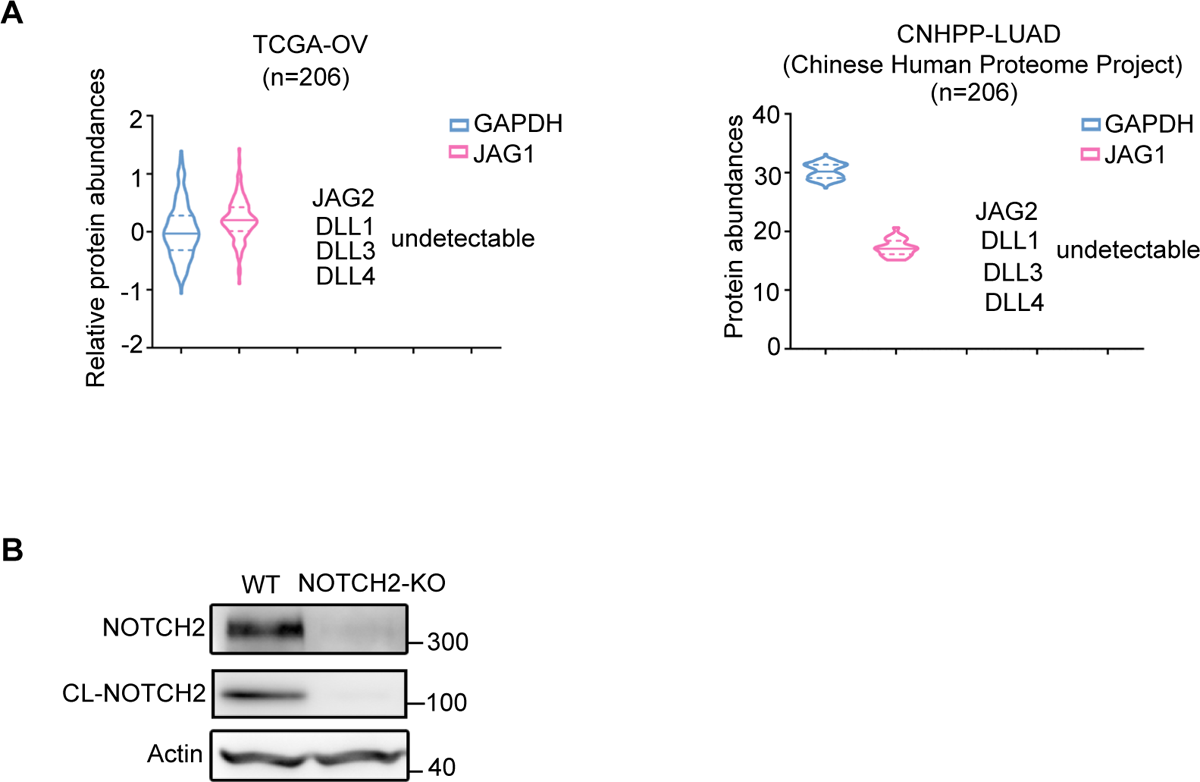
Activation of NOTCH2 signaling confers resistance to taxol-induced cell death. **(A)** Analysis of protein levels of NOTCH ligands in ovarian and lung cancer using TCGA and CNHPP data. **(B)** Western blot analysis of NOTCH2 knockout efficiency in A549 cells.

**Figure S3.**
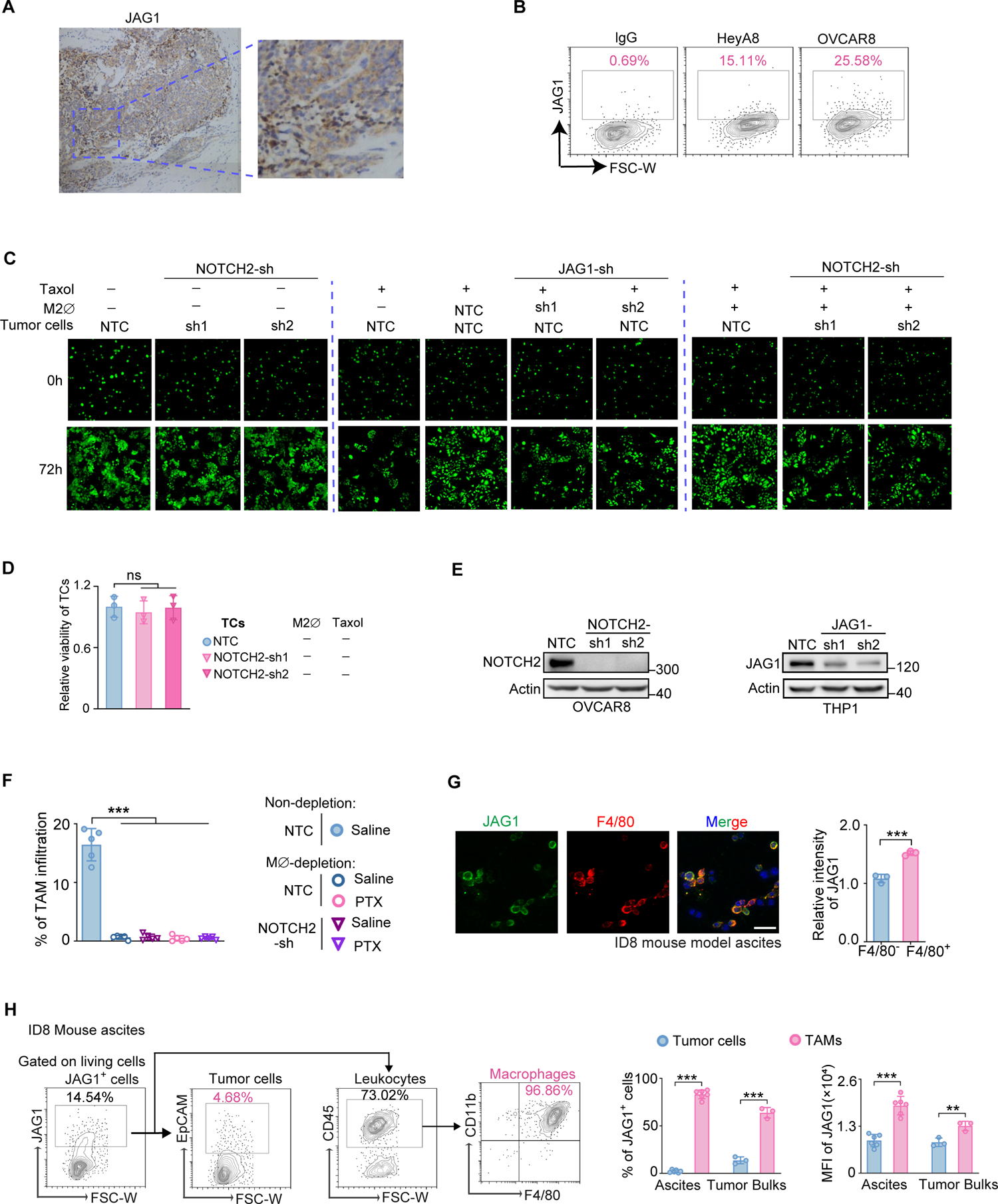
JAG1 expressed on the macrophages is primarily responsible for activating NOTCH2 signaling and promotes tumor cells resistance to taxol. **(A)** Immunohistochemical staining of JAG1 in tumor section from recurrent ovarian cancer patients. Framed image was enlarged on the right. **(B)** JAG1 expression in cancer cell lines were analyzed by FACS. **(C)** Representative images of OVCAR8-GFP cells alone or in co-culture with THP-1-derived M2 macrophages (M2ϕ). **(D)** Quantification of the relative viability of GFP^+^ cells culture is shown. **(E)** Western blot analysis of the knockdown efficiency of NOTCH2 and JAG1 in OVCAR8 and THP-1 cells, respectively. **(F)** Percentage of infiltrated TAM in ID8 mouse ascites in figure 3E. **(G)** Immunofluorescence and the quantitation of JAG1 expressed on macrophages in the ascites of ID8 tumors. Three ascites were analyzed for each group. Scale bars, 10 μm. **(H)** FACS analysis was conducted to assess the expression of JAG1 in TAM from the ascites of ID8 mice. Data are shown as mean ±SD. Data of (D), (G) and (H) were obtained from three experimental replicates. p values were determined by two-tailed Student’s t test (**p < 0.01, ***p < 0.001).

**Figure S4.**
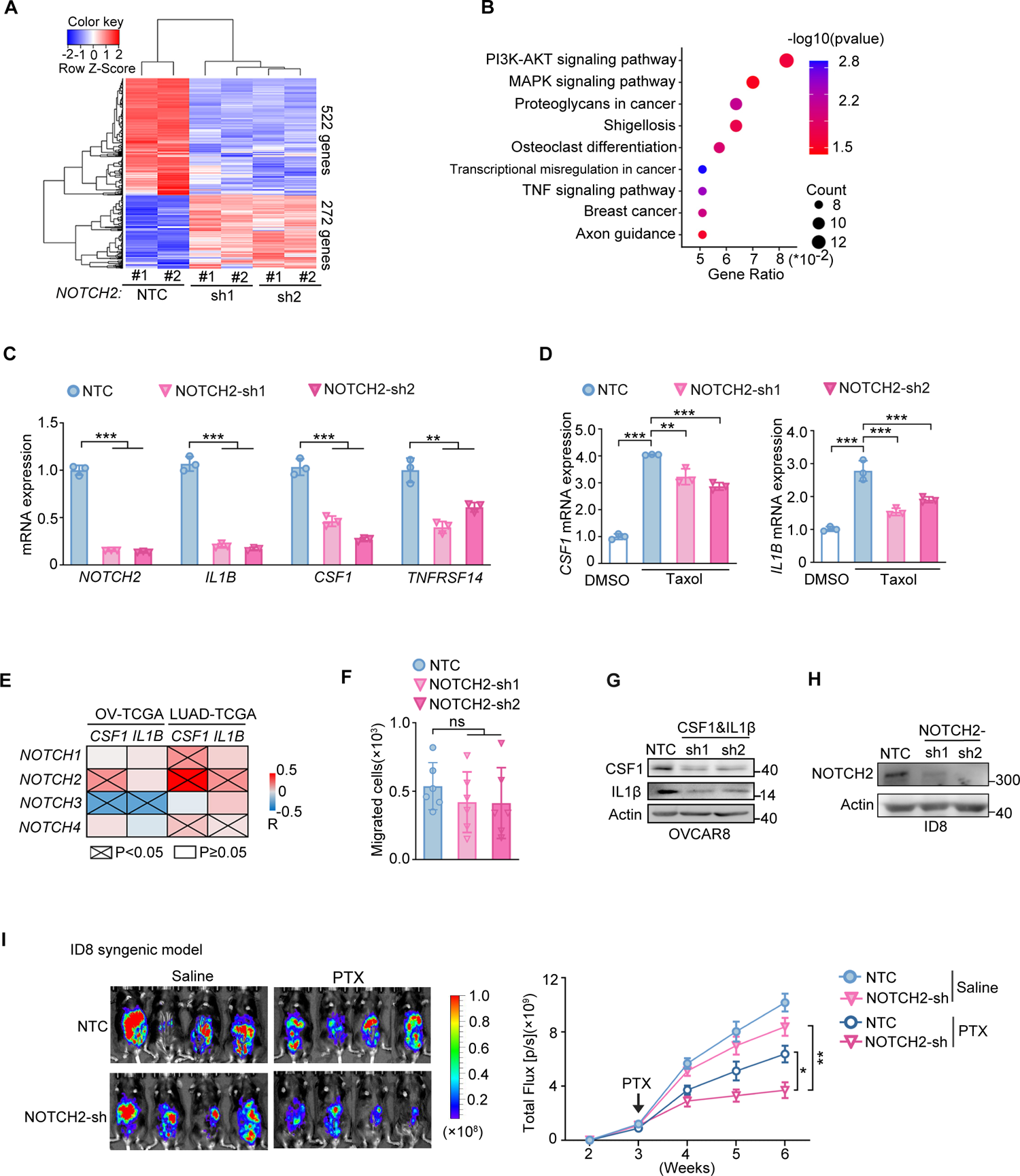
NOTCH2 upregulation activates pro-survival pathways and stimulates the recruitment of pro-tumor macrophages. **(A)** Heatmap illustrates differentially expressed genes (DEGs) in OVCAR8 cells transfected with NOTCH2-shRNA or non-target control (NTC) vector. **(B)** Kyoto Encyclopedia of Genes and Genomes (KEGG) pathway analysis of DEGs in RNA-Seq data (NOTCH2 knock-down vs Non-target control (NTC)). **(C)** RT– PCR detection of the DEG mRNA levels in OVCAR8 cells after NOTCH2 knockdown. **(D)** RT–PCR analysis of *CSF1* and *IL1B* mRNA levels in A549 cells after NOTCH2 knockdown. Cells were treated with DMSO or 100 nM taxol for 48 hrs. **(E)** Pearson correlation between NOTCH receptors and CSF1 and IL1B mRNA expression in ovarian cancer and lung adenocarcinoma of NSCLC from the TCGA database. **(F)** Chemotactic with DMSO. **(G)** Knockdown level of CSF1 and IL-1B in OVCAR8 cells. **(H)** Knockdown level of NOTCH2 in ID8 cells. **(I)**. Representative images of mice carrying ID8-luciferase cells transfected with inducible NOTCH2 knockdown shRNA three weeks after treatment (left panel) and the corresponding tumor growth curves (right panel). Data are shown as mean ±SD, and data of (C), (D) and (F) were obtained from three experimental replicates. p values were determined by two-tailed Student’s t test (ns, not significant; **p < 0.01, ***p < 0.001).

**Figure S5.**
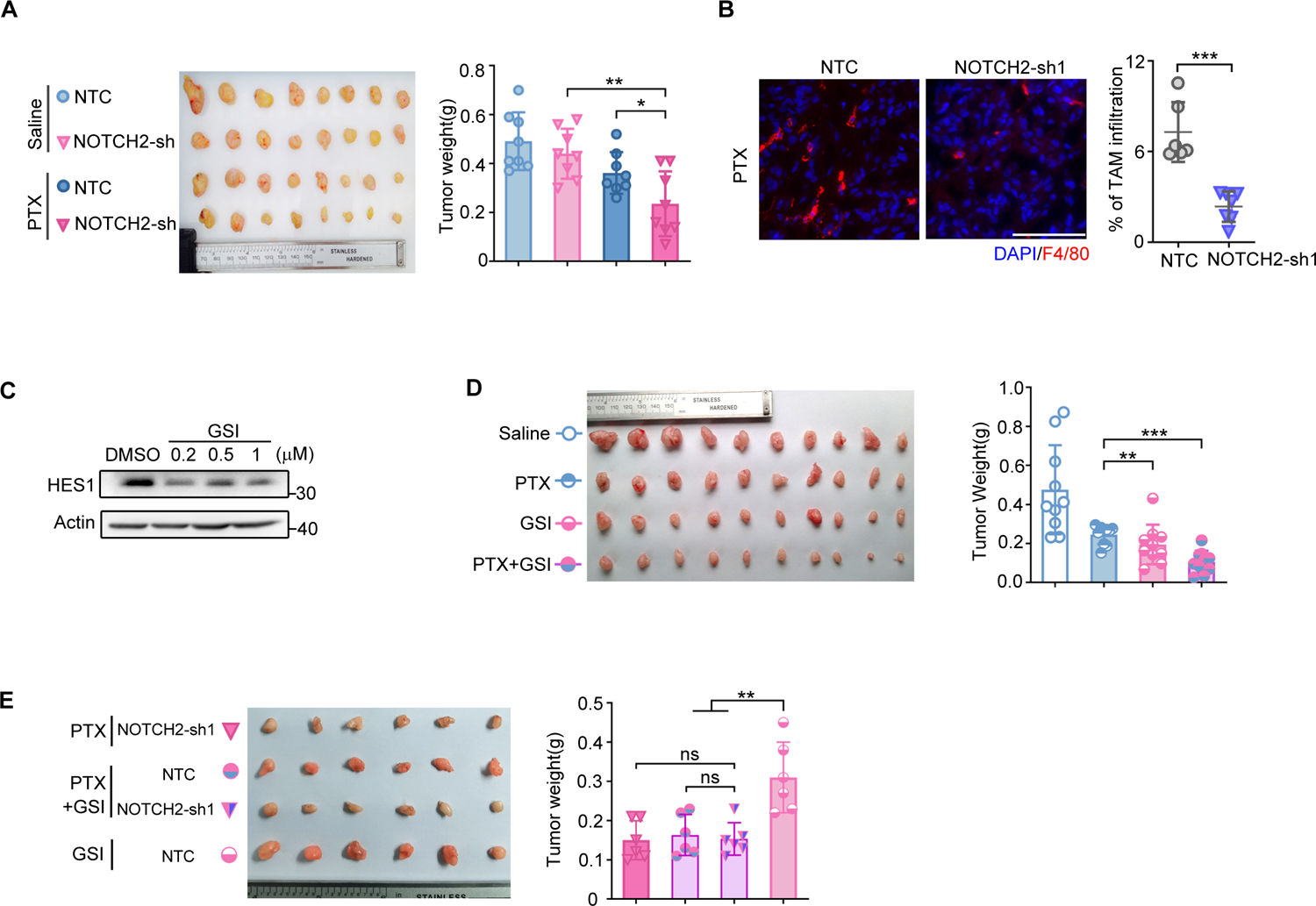
NOTCH2 inhibition sensitizes tumor response to paclitaxel in multiple mouse tumor models. **(A)** Representative dissected tumors (left) and tumor weight (right panel) of A549-xenograft mice. A549 cells with doxycycline inducible NOTCH2 knockdown (sh) or non-target control (NTC) were injected and 10 mg/kg paclitaxel was administered by intraperitoneal injection when tumor volume reached about 100 mm^3^. Error bars are SEM. **(B)** Representative images of tumor sections from OVCAR8-xenograft mice stained with F4/80 and DAPI. Quantification of the percentage of F4/80+ macrophage cells is shown. Scale bars, 20 μm. **(C)**Western blot analysis of GSI effect on inhibiting NOTCH signaling. **(D)** Tumor growth of xenograft mice carrying A549 cells treated with GSI and paclitaxel either alone or in combination. Representative tumors (left panel) and tumor weight (right panel) for each group are shown. **(E)** Mice carrying OVCAR8 wildtype or NOTCH2-knockout xenografts were treated with GSI and paclitaxel. p values were determined by two-tailed Student’s t test and log-rank test (ns, not significant; **p < 0.01, ***p < 0.001).

**Figure S6.**
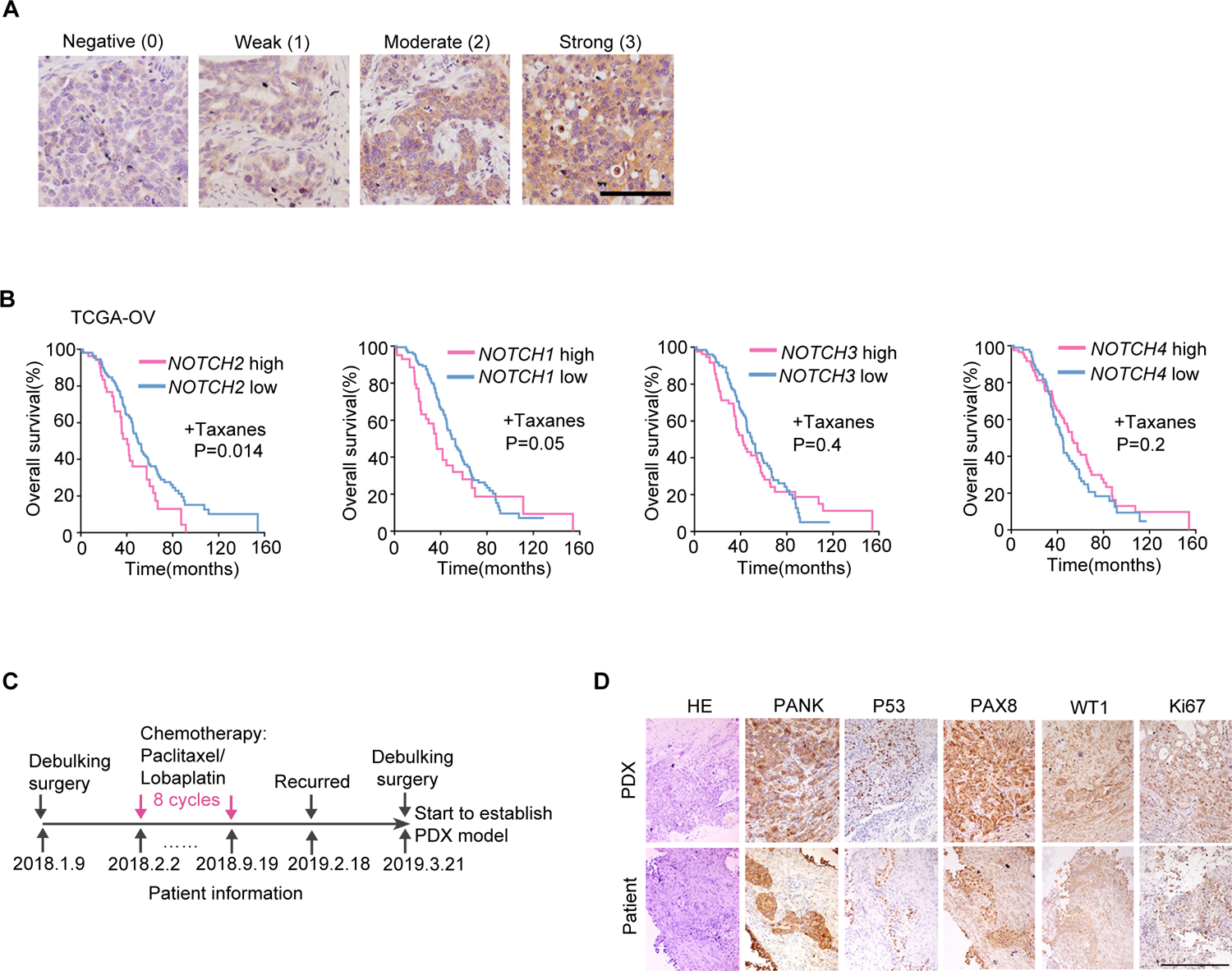
High NOTCH2 level is associated with chemoresistance of clinical samples. **(A)** Representative IHC staining images of tumor specimens from ovarian cancer patients. Staining intensity of 0 (negative), 1 (weak), 2 (moderate), and 3 (strong) of NOTCH2 are shown. **(B)** Survival curves of ovarian cancer patients from the TCGA database stratified according to treatment (taxanes versus others) and expression levels of NOTCH1, 2, 3, 4 (cutoff: auto-select best cutoff). **(C)** Information of the ovarian cancer patient whose tumor tissue was used to establish the PDX mouse model. **(D)** Immunohistochemistry (IHC) detection of ovarian cancer markers in tissues from the patient and PDX model. p values were determined by the log-rank test.

**Table S1.** Patients’ information.

**Table S2.** Reagent and Resource.

## Notes

### Competing Interest Statement

The authors have declared no competing interest.

