## Supplemental Table 1 for "Targeting NOTCH2-JAG1 juxtacrine signaling reverses macrophage-mediated tumor resistance to taxol"

### Table S1

| Reagent and Resource | Source | Identifer |
|---|---|---|
| Reagent | | |
| Red Blood Cell Lysis Buffer | Beyotime Biotechnology | #C3702 |
| Antibodies | | |
| CD68 (D4B9C) XP® Rabbit mAb | Cell signal Technology | #76437 |
| HES1 (D6P2U) Rabbit mAb | Cell signal Technology | #11988 |
| Mouse Monoclonal Anti-Actin | Proteintech | #66009-1-Ig |
| Mouse monoclonal anti-Bcl2 | BD biosciences | #610538 |
| Mouse Monoclonal Anti-b-Tubulin | Sigma-Aldrich | #T4026 |
| Mouse monoclonal anti-F4/80 | BioLegend | #123101 |
| Mouse Monoclonal Anti-GAPDH | Proteintech | #60004-1-Ig |
| Mouse monoclonal anti-Jagged1 | Santa Cruz Biotechnology | #sc-390177 |
| Mouse Monoclonal Anti-Ki67 | Leica Biosystems | #ACK02 |
| Mouse monoclonal anti-p38 MAPK | Proteintech | #66234-1-Ig |
| Mouse monoclonal anti-p53 | Santa Cruz Biotechnology | #sc-126 |
| Mouse monoclonal anti-PAX8 | Proteintech | #60145-4-Ig |
| Notch1 (D1E11) XP® Rabbit mAb | Cell signal Technology | #4851 |
| Notch2 (D67C8) XP® Rabbit mAb | Cell signal Technology | #4530 |
| Phospho-p38 MAPK (Thr180/Tyr182) (D3F9) XP® Rabbit mAb | Cell signal Technology | #4511 |
| Rabbit Caspase-3 Antibody | Cell signal Technology | #9662 |
| Rabbit polyclonal anti-CPEB1 | Proteintech | #13274-1-AP |
| Rabbit polyclonal anti-Cyclin-B1 | Proteintech | #55004-1-AP |
| Rabbit polyclonal anti-Mcl1 | Santa Cruz Biotechnology | #sc-819 |
| Rabbit polyclonal anti-Notch3 | Proteintech | #55114-1-AP |
| Rabbit polyclonal anti-Notch4 | Abclonal | #A8303 |
| Rabbit polyclonal anti-pan-keratin | Proteintech | #26411-1-AP |
| Rabbit polyclonal anti-PARP1 | Proteintech | #13371-1-AP |
| Rabbit polyclonal anti-WT1 | Proteintech | #12609-1-AP |
| Purified anti-mouse CD16/32 Antibody | BioLegend | #101302 |
| APC/Cy7 anti-mouse CD45.2 Antibody | BioLegend | #109824 |
| APC anti-mouse/human CD11b Antibody | BioLegend | #101212 |
| PE anti-mouse F4/80 Antibody | BioLegend | #123109 |
| PE/Cy7 anti-human CD206 | BioLegend | #321123 |
| Biological Samples | | |
| Ovarian cancer samples | The First Affiliated Hospital of the University of Science and Technology of China | This paper |
| Chemicals, Peptides, and Recombinant Proteins | | |
| BI2536 | Selleckchem | #S1109 |
| Carboplatin | Selleckchem | #S1215 |
| Cordycepin | Sigma | #C9137 |
| Cycloheximide | Sigma | #239763 |
| D-Luciferin | GoldBio | #115144-35-9 |
| Dimethyl sulfoxide (DMSO) | Sangon Biotech | #A600163-0250 |
| Nocodazole | Sigma | #M1404 |
| Paclitaxel | Sigma-Aldrich | #1491332 |
| Polybrene | Sigma-Aldrich | #TR-1003 |
| Puromycin | Selleckchem | #S7417 |
| RO4929097 | Selleckchem | #S1575 |
| SB203580 | Selleckchem | #S1076 |
| STLC | Sigma-Aldrich | #SPE006 |
| Thymidine | Sigma | #T1895 |
| Human recombinant CSF1 | R&D Systems | #216-MC-010 |
| Human recombinant IL1B | R&D Systems | #201-LB-010 |
| Human recombinant IL13 | R&D Systems | #213-ILB-025 |
| Human recombinant IL4 | R&D Systems | #204-IL-020 |
| Critical Commercial Assays | | |
| Bradford Protein Assay Kit | Sangon Biotech | # C503031 |
| CellTiter-Glo® Luminescent Cell Viability Assa | Promega | #G7571 |
| Human IL-1 beta ELISA Kit | Abclonal | #RK00001 |
| Human M-CSF ELISA Kit | Abclonal | #RK00044 |
| Depoisted Data | | |
| RNA-seq(Ribosome profiling) | GEO | GSE158569 |
| RNA-seq(Notch2 Knockdown) | GEO | GSE158569 |
| Experimental Models: Cell Lines | | |
| HEK 293T | ATCC | N/A |
| HeLa | ATCC | N/A |
| OVCAR8 | ATCC | N/A |
| HeyA8 | ATCC | N/A |
| ID8 | N/A | Kindly provided by Dr. K.F. Roby |
| THP-1 | ATCC | N/A |
| OVCAR8-Gipz-RH4346 Stable cell line | This paper | N/A |
| OVCAR8-Gipz-Notch2-sh1 Stable cell line | This paper | N/A |
| OVCAR8-Gipz-Notch2-sh2 Stable cell line | This paper | N/A |
| OVCAR8-Tripz-RH4346 Stable cell line | This paper | N/A |
| OVCAR8-Tripz-Notch2-sh1 Stable cell line | This paper | N/A |
| OVCAR8-Tripz-Notch2-sh2 Stable cell line | This paper | N/A |
| HeyA8-Gipz-RH4346 Stable cell line | This paper | N/A |
| HeyA8-Gipz-Notch2-sh1 Stable cell line | This paper | N/A |
| HeyA8-Gipz-Notch2-sh2 Stable cell line | This paper | N/A |
| ID8-Tripz-RH4346 Stable cell line | This paper | N/A |
| ID8-Tripz-Notch2-sh1 Stable cell line | This paper | N/A |
| ID8-Tripz-Notch2-sh2 Stable cell line | This paper | N/A |
| THP-1-PLKO.1 Stable cell line | This paper | N/A |
| THP-1-PLKO.1-JAG1-sh1 Stable cell line | This paper | N/A |
| THP-1-PLKO.1-JAG1-sh2 Stable cell line | This paper | N/A |
| Experimental Models: Organisms/Strains | | |
| NOD SCID | Charles River | N/A |
| C57BL/6JNifdc | SLAC Laboratory Animal | N/A |
| Balb/c nude | SLAC Laboratory Animal | N/A |
| Ovarian cancer PDX model | This paper | N/A |
| Oligonucleotides | | |
| qRTPCR primer for human specific Notch2 | This paper | N/A |
| Forward-5'-CCTTCCACTGTGAGTGTCTG/ | | |
| Reverse-5'-AGGTAGCATCATTCTGGCAG( | | |
| qRTPCR primer for human specific ITGB4 | This paper | N/A |
| Forward-5'-GCAGCTTCCAAATCACAGAGG | | |
