## Supplemental Table 2 for "Targeting NOTCH2-JAG1 juxtacrine signaling reverses macrophage-mediated tumor resistance to taxol"

### Table S2

| Number | Age | Histologic subtypes | Neo/adjuvant chemotherapy | Chemotherapy regimens | Chemotherapy Cycles | PFS(months) | Notch2(H-Score) Primary | Recurrent |
|---|---|---|---|---|---|---|---|---|
| 1 | 55 | HGSO-IIIC | Adjuvant | Taxane and platinum | 9 | 12 | 0.8 | 3.5 |
| 2 | 67 | HGSO-IV | Neo | Taxane and platinum | 3 | none | 2.0 | 4.0 |
| 3 | 51 | HGSO-IIIC | Adjuvant | Taxane and platinum | 9 | 5 | 0.0 | 10.0 |
| 4 | 66 | HGSO-IV | Adjuvant | Taxane and platinum | 8 | 15 | 0.2 | 4.0 |
| 5 | 46 | HGSO-IIIC | Neo | Taxane and platinum | 4 | none | 2.0 | 1.6 |
| 6 | 53 | HGSO-IV | Neo | Taxane and platinum | 3 | none | 0.7 | 3.0 |
| 7 | 65 | Clear cell carcinoma-IV | Adjuvant | Taxane and platinum | 4 | 7 | 3.5 | 8.0 |
| 8 | 47 | HGSO-IIIC | Adjuvant | Taxane and platinum | 8 | 14 | 2.3 | 9.3 |
| 9 | 60 | poorly differentiated Adnocarcinoma IV | Neo | Taxane and platinum | 4 | none | 8.0 | 5.0 |
| 10 | 62 | HGSO-IIIC | Adjuvant | Taxane and platinum | 13 | 14 | 3.0 | 5.3 |
| 11 | 61 | HGSO-IV | Adjuvant | Taxane and platinum | 3 | 12 | 0.3 | 6.0 |
| 12 | 57 | Serous carcinoma | Adjuvant | Taxane and platinum | 3 | 2 | 3.0 | 8.7 |
| 13 | 61 | HGSO | Neo | Taxane and platinum | 9 | none | 3.5 | 4.8 |
| 14 | 66 | HGSO-IIIC | Adjuvant | Taxane and platinum | 9 | 11 | 2.3 | 8.6 |
| 15 | 55 | HGSO IIIC | Adjuvant | Taxane and platinum | 15 | 23 | 0.3 | 5.3 |
| 16 | 52 | HGSO IIIC | Adjuvant | Taxane and platinum | 24 | 64 | 0.0 | 2.0 |
| 17 | 54 | Serous carcinoma IIC | Adjuvant | Taxane and platinum | 7 | 43 | 0.3 | 10.0 |
| 18 | 61 | Serous carcinoma IIC | Adjuvant | Taxane and platinum | 6 | 59 | 0.2 | 4.0 |
| 19 | 49 | HGSO-IIIC | Adjuvant | Taxane and platinum | 11 | 30 | 3.0 | 4.7 |
| 20 | 45 | HGSO-IIIC | Adjuvant | Taxane and platinum | 8 | 14 | 4.80 | 6.40 |
| 21 | 65 | ovarian sex cord stromal tumor | Adjuvant | Taxane and platinum | 1 | 4 | 0.00 | 4.00 |
| 22 | 49 | Poorly differentiated Adnocarcinoma IIIC | Neo | Taxane and platinum | 3 | none | 0.00 | 4.00 |
| 23 | 59 | Poorly differentiated Adnocarcinoma IIIC | Adjuvant | Taxane and platinum | 3 | 2 | 3.20 | 5.33 |
| 24 | 48 | Poorly differentiated Adnocarcinoma IIIC | Adjuvant | Taxane and platinum | 2 | 2 | 1.60 | 6.40 |
| 25 | 63 | Poorly differentiated Adnocarcinoma IIIC | Adjuvant | Taxane and platinum | 3 | 20 | 1.60 | 3.00 |
| 26 | 56 | Poorly differentiated Adnocarcinoma IIIC | Adjuvant | Taxane and platinum | 20 | 57 | 2.40 | 9.60 |
| 27 | 52 | Poorly differentiated Adnocarcinoma IV | Neo | Taxane and platinum | 3 | none | 4.00 | 0.00 |
| 28 | 54 | HGSO-IIIC | Adjuvant | Taxane and platinum | 13 | 27 | 3.20 | 2.00 |

Age at time of surgery.
The FIGO staging system
Progression free survival (months), The time from First chemotherapy to tumor recure.
